# Loss of PTPRB function remodels VEGFR1 activation in tumors overexpressing the receptor tyrosine kinase

**DOI:** 10.64898/2026.08.17.745152

**Authors:** Sayar Ghosh, Abheek Pathak, Aayushee Ghosh, Manas Pratim Chakraborty, Bikram Das, Saptarshi Pyne, Rahul Das

## Abstract

The VEGF Receptor-1 (VEGFR1) is a deceptive receptor tyrosine kinase (RTK). In early embryonic development, VEGFR1 negatively regulates angiogenesis by acting like a decoy receptor. Ligand binding transiently phosphorylates the receptor and induces a weak activation, even at high receptor density. Yet, in multiple cancers, overexpression of VEGFR1 plays a central role in tumor vascularization and growth. Unlike many pro-oncogenic RTKs, extensive patient data analysis revealed no somatic mutation in VEGFR1 that may spontaneously activate the tyrosine kinase. The mechanism by which VEGFR1 is activated in cancers has remained an open question for more than two decades. Here, we evaluated the multi-omics profiles of VEGFR1 and its regulators in a pan-cancer database. We observed an inverse correlation between VEGFR1 and PTPRB phosphatase expression in KIRC patients and disease outcome. We observed that patients overexpressing VEGFR1 and deficient in PTPRB expression have a lower likelihood of survival. Using super-resolution single-cell imaging, we discovered that inhibiting PTPRB spontaneously activates VEGFR1 by inducing ligand-independent dimerization, possibly by shifting the equilibrium toward the active state. PTPRB inhibition induces sustained, ligand-dependent phosphorylation of VEGFR1, which may promote tumor vascularization. We conclude that a subtle phosphatase imbalance is fundamental in determining VEGFR1’s role in pathological angiogenesis in tumors.

## Introduction

The vascular endothelial growth factor receptor-1 (VEGFR1) is an inefficient and elusive receptor tyrosine kinase (RTK) compared with its family member, VEGFR2^1–4^. During early embryogenesis, VEGFR1 negatively regulates the binding of proangiogenic ligand, VEGF-A, to the main angiogenic receptor VEGFR2^2,5^. VEGFR1 competes for the ligand with a ten-fold higher affinity^6,7^. Ligand binding to VEGFR1 induces a transient and weak tyrosine autophosphorylation^4,8,9^. The low-amplitude signal thus generated does not induce cell proliferation, migration, or tube formation in endothelial cells, vascular smooth muscle cells, or fibroblasts^6,8,10,11^. Suggesting tyrosine kinase activity of VEGFR1 is not required for angiogenesis. However, VEGFR1 kinase activation is necessary for monocyte and macrophage migration^6,12^.

Ligand-independent activation of RTKs, such as EGFR, FGFR, and VEGFR2, at high receptor density is often linked to the pathology of multiple cancers^13–18^. On the contrary, VEGFR1, when overexpressed in macrophage cells, does not exhibit ligand-independent tyrosine phosphorylation^9^. Despite that, VEGFR1 signaling has emerged as a crucial factor regulating pathological angiogenesis, tumor development, and survival^19–23^. The VEGFR1-expressing bone marrow-derived myeloid cells play an important role in initiating tumor growth^21^. Metastasis-associated macrophages expressing VEGFR1 support tumor seeding and facilitate cancer metastasis^24,25^. Where VEGFR1 links the ligand binding to the colony-stimulating factor 1 (Csf1) mediated inflammatory response, promoting metastasis in breast cancer. Impaired VEGFR1 signaling in the nude mouse xenograft model resulted in a significant reduction in tumor vascularization and growth in kidney renal clear cell carcinoma (KIRC)^26^ and glioma^27^. Thus, VEGFR1 and its ligands have emerged as potential drug targets for antiangiogenic therapy^22^. Blocking VEGFR1 activation using a monoclonal antibody^28^ inhibits the metastasis of multiple tumor models and reduces cancer-related pain in pancreatic ductal adenocarcinoma^29^. Yet, it remains unclear how overexpressing VEGFR1 in tumor cells activates the kinase.

The structure of VEGFR1 comprises an extracellular ligand-binding domain (ECD), connected to a cytosolic tyrosine kinase domain (KD) (Figure 1)^30,31^. A single-pass transmembrane (TM) segment and a cytosolic juxtamembrane (JM) segment connect the ECD to the catalytic unit. The C-terminal tail following the KD has multiple tyrosine residues that serve as an adaptor module upon phosphorylation. The VEGFR1 predominantly exists as a monomer in the autoinhibited (inactive) conformation (Figure 1b)^32,33^. The monomeric state is maintained by the electrostatic repulsion between the immunoglobulin domains of ECD^34,35^. The autoinhibited conformation of the KD is stabilized by the PDGFR-like JM-in conformation, where Y794 occupies the catalytic site as a pseudosubstrate^36,37^. An additional electrostatic interaction between the negatively charged residues in the JM-S segment and the positively charged patch on the C-lobe of the KD further strengthens the JM inhibitory interactions^9^. The additional JM autoinhibitory interactions are a unique feature of VEGFR1 and are absent from the rest of the VEGFR family. The strong JM autoinhibition is enough to prevent ligand-independent activation of VEGFR1 at high receptor density (Figure 1b). As a general mechanism, ligand binding induces a conformational rearrangement of the ECD, thereby allowing receptor dimerization (Figure 1b)^18,38–40^. The conformational rearrangement at the ECD releases the JM autoinhibition by reorienting the JM segment into an out conformation^36,37^. The slow release of the JM autoinhibition, in turn, imparts a transient tyrosine phosphorylation in the C-terminal tail^9^. It is perplexing to find that in multiple VEGFR1-overexpressing cancers, JM autoinhibition is spontaneously released. The mechanism of which is still unknown.

**Figure 1:**
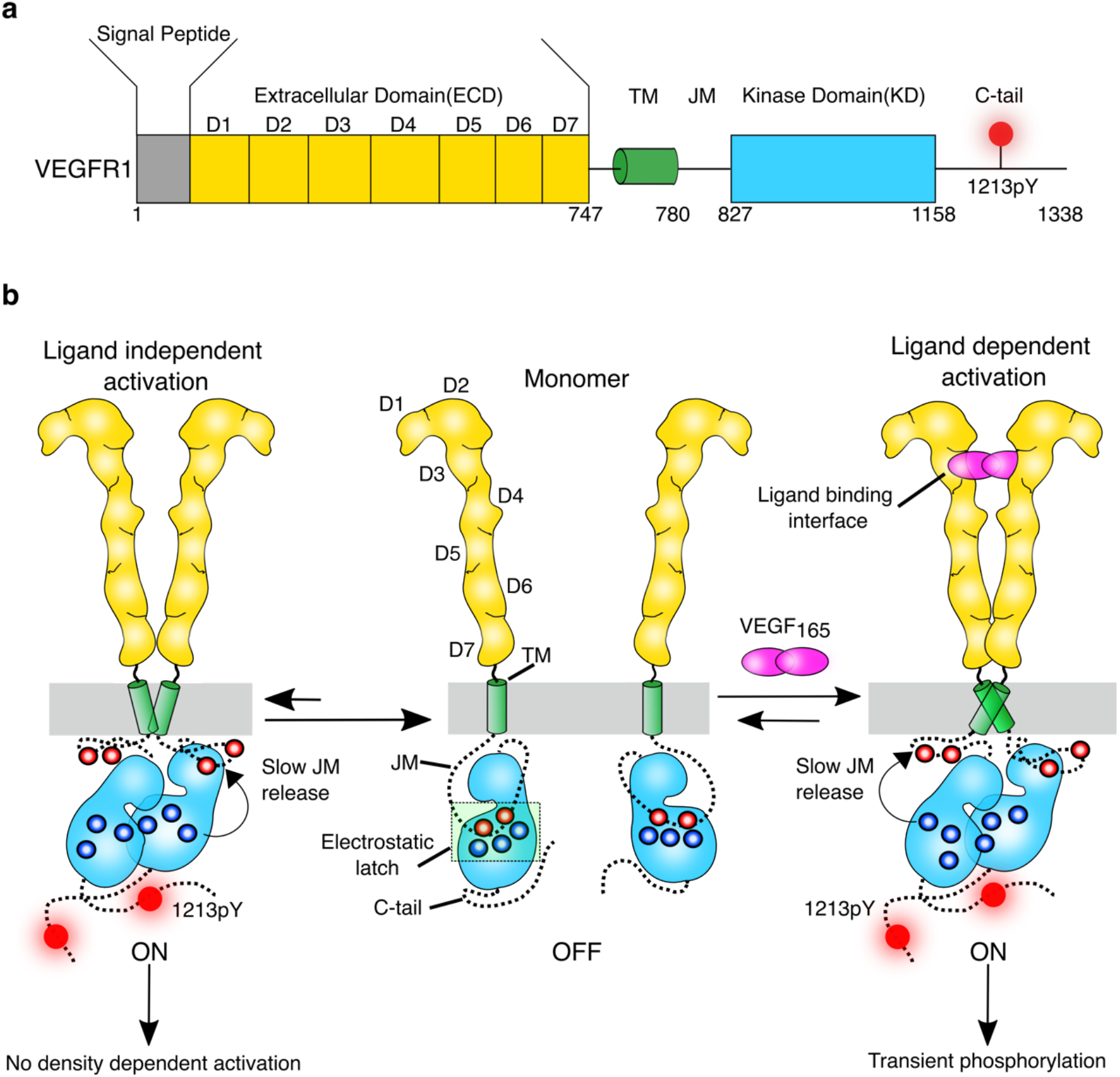
Model for VEGFR1 activation. **a)** Schematic representation of the domain architecture of VEGFR1. The extracellular domain (ECD), transmembrane segment (TM), juxtamembrane segment (JM), kinase domain (KD), and C-terminal tail (C-tail) are labeled. The phosphor-tyrosine residue (1213pY) in the C-tail used to monitor VEGFR1 autophosphorylation is highlighted. **b)** Cartoon representation of ligand-independent and ligand-dependent activation of VEGFR1. The electrostatic latch that stabilizes the autoinhibited conformation is labeled. The electrostatic interactions between the positive charge residues (blue circle) in the C-lobe of the kinase domain and the negatively charged residues (red circle) in the JM segment are highlighted. The relative arrow length indicates the preferred direction of structural rearrangement. The schematics are designed using Inkscape Ver1.4.3.

In this study, we extensively examine the genomic, transcriptomic, and phosphorylation profiles of patient data from The Cancer Genome Atlas (TCGA) overexpressing VEGFR1. We observed that VEGFR1 is significantly overexpressed in KIRC, also known as clear cell Renal cell carcinoma (ccRcc). A markedly elevated level of VEGFR1 phosphorylation was observed in KIRC samples, suggesting that the receptor may be persistently phosphorylated. Counterintuitively, in the COSMIC (Catalog of Somatic Mutations in Cancer) repository, we could not find any high-frequency mutation in VEGFR1 that might explain its sustained activation in cancer. Using KIRC as a model, we explore the gene regulatory network (GRN) of VEGFR1 in the tumor. GRN analysis suggests a possible link between reduced expression of protein tyrosine phosphatase receptor type B (PTPRB) and sustained phosphorylation of VEGFR1. We observed that loss of PTPRB expression correlates with lower survival among KIRC patients. Finally, using single-cell imaging, we presented a mechanism explaining how blocking tyrosine phosphatases releases JM inhibition and improves the phosphotyrosine half-life (t_1/2_) of VEGFR1 at the membrane. We observed that inhibiting PTPRB induces ligand-independent activation of VEGFR1 and sustains ligand-independent phosphorylation by stabilizing an active dimer.

## Results

### VEGFR1 is Overexpressed in KIRC

We begin with a comprehensive analysis of TCGA cancer patient data to assess the transcriptomic profiles of VEGFR1, VEGFR2, and VEGF-A in pan-cancer data (Figures 2a, S1a-b)^41^. For our study, we consider mRNA profiles from tumor and adjacent normal tissues, with sample sizes greater than 50. We observed that in COADREAD (colonic and rectal adenocarcinoma) and KIRC tissues, VEGFR1 is significantly overexpressed compared with adjacent normal tissues (Figure 2a). We also noted overexpression of VEGFR2 in KIRC tissue (Figure S1a), which correlates with the overexpression of its ligand VEGF-A (Figure S1b). Previous independent multi-omics studies have extensively evaluated genomic, epigenomic, transcriptomic, proteomic, and phosphoproteomic data, revealing that VEGFR1 may be overexpressed in KIRC^42,43^. Deregulation of von Hippel-Lindau (VHL) is found in almost all KIRC tumors, leading to accumulation of hypoxia-inducible factors (HIF)^43–45^. This, in turn, promotes tumor vascularization by increasing transcription of VEGFR1 and its ligand^21,46,47^. Independent studies have shown that knocking down VEGFR1 in the KIRC cell line^26^ or sequestering the VEGFR1 ligand with monoclonal antibodies^28^ inhibits tumor growth and metastasis. Together, these findings imply that overexpression of VEGFR1 plays an important role in the pathological angiogenesis in KIRC. However, the fundamental question of how VEGFR1 activation is differently regulated in the tumor tissues overexpressing the receptor remained unanswered. Since the multi-omics profile of KIRC has been exhaustively characterized,^42,43^ we use KIRC as a model system for patient data analysis in our current study.

**Figure 2:**
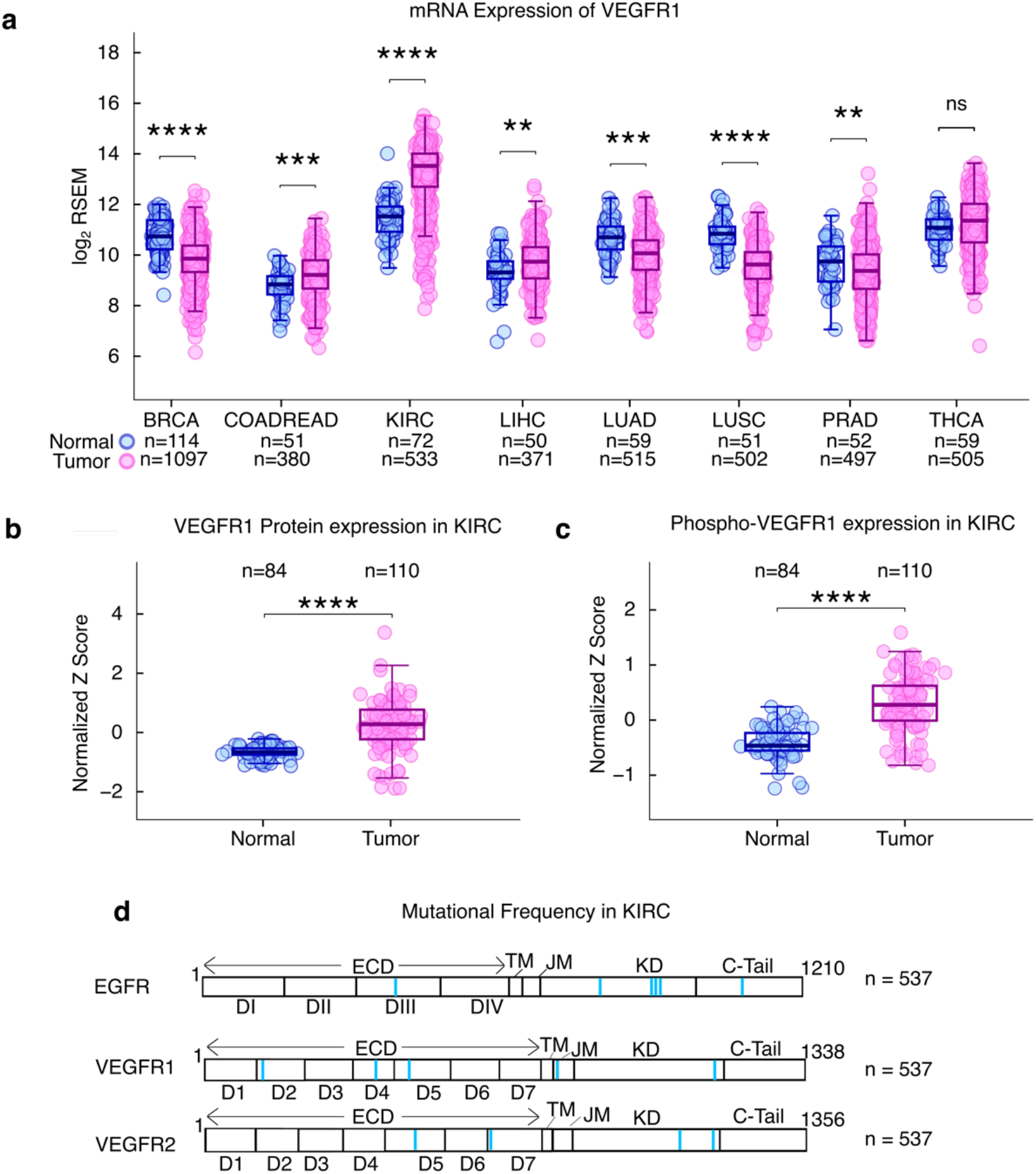
VEGFR1 expression and activation in KIRC patients. **a)** Plot of normalized mRNA level of VEGFR1 (log_2_RSEM) expressed in normal tissue (blue) and tumor tissue (pink) from the indicated cancer patient. The patient information was retrieved from the TCGA-curated cancer dataset. **b)** Plot of normalized VEGFR1 expression in normal (blue) and tumor (pink) tissue from KIRC patients. **c)** Plot of normalized VEGFR1 phosphorylation level in normal (blue) and tumor tissue (pink) in KIRC patients. **d)** Mutation frequency in EGFR, VEGFR1, and VEGFR2 observed in KIRC patients is mapped on the secondary structure of the receptor. The region where no mutations were detected is shown in white. The blue vertical line indicates the number of patients carrying mutations in fewer than 5 of 537 samples. RSEM-RNA-Seq by Expectation Maximization, BRCA-Breast Cancer, COADREAD-Colonic and rectal adenocarcinoma, KIRC-Kidney renal clear cell carcinoma, LIHC-Liver hepatocellular carcinoma, LUAD-Lung adenocarcinoma, LUSC-lung squamous cell carcinoma, PRAD-Prostate adenocarcinoma, THCA-Thyroid carcinoma In panels a-c, each circle represents a patient. The solid horizontal line is the median, and the error indicates ±SD. Comparison between the two groups was made using Student’s t-test. Statistical significance was assessed using the following criteria: \**p*< 0.05, \*\**p*< 0.01, \*\*\**p*< 0.001, \*\*\*\**p*< 0.0001, and ns denotes not significant. **d)** The schematic is made using Inkscape Ver1.4.3.

### VEGFR1 shows sustained Phosphorylation in KIRC

VEGFR1 is difficult to activate in cell-based studies^4,8,48,49^, which undergoes transient autophosphorylation after ligand binding^9^. In contrast, VEGFR2 shows sustained tyrosine phosphorylation upon VEGF-A binding^9^. Therefore, we next investigate whether overexpression of VEGFR1 remodels the phosphorylation level of the receptor in KIRC tissue and compare that with VEGFR2 phosphorylation (Figures 2 b-c and S1 c-d). The proteomics and phosphoproteomics data for VEGFR1 and VEGFR2 in KIRC tissues and normal adjacent tissues were obtained from the Clinical Proteomic Tumor Analysis Consortium (CPTAC)^43^. For VEGFR1, we observed that the transcriptomic profile correlates with the proteomic profile (Figure 2a-b). The overexpression of VEGFR1 also leads to hyperphosphorylation of tyrosine residues of VEGFR1 in KIRC tissues compared to normal adjacent tissues (Figure 2c). Suggesting that the VEGFR1 overexpression induces a sustained tyrosine phosphorylation in the tumor. We noted a marginal increase in VEGFR2 expression in KIRC tissue (Figure S1c). As anticipated, VEGFR2 showed a sustained tyrosine phosphorylation (Figure S1d). It may be noted that VEGFR1 is poorly phosphorylated upon ligand binding in non-cancerous cells^8,11^. Multiple factors determine the phosphotyrosine status of RTKs at the plasma membrane, including mutations, increased receptor density, or changes in the phosphatase profile of tumor cells. Therefore, we next examine the factors that may induce spontaneous (ligand-independent) and sustained (upon ligand binding) phosphorylation of VEGFR1 in KIRC.

### No spontaneously activating VEGFR1 mutant was detected in KIRC

Oncogenic mutations in RTKs are often associated with cancer development and metastasis^50^. For example, the Q472H mutation, which spontaneously activates VEGFR2, is frequently mutated in melanoma^51^. In EGFR, single amino acid substitutions in the extracellular domain (A289V) or kinase domain (L858R) are linked to glioblastoma^52^ or non-small cell lung carcinoma^53^, respectively.

The autoinhibited structure of the VEGFR1 is stabilized by the repulsive interaction between the ECD^35,40^ and by JM inhibition^9,54^. Mutating N1050D in the activation loop^55^ or replacing either the JM^9,56^ or the C-terminal tail with that of VEGFR2 activates the kinase^55^. We recently showed that a destabilizing mutation in the JM-latch, along with the simultaneous removal of Y794 from the catalytic pocket, induced ligand-independent activation of VEGFR1 at high receptor density^9^. The mutations also remodel transient VEGFR1 tyrosine phosphorylation into sustained phosphorylation. We therefore mined the COSMIC (Catalog of Somatic Mutations in Cancer)^57^ data for mutations that might spontaneously activate VEGFR1 and compared them with those in VEGFR2 and EGFR. In the COSMIC pan-cancer database, we identified frequently occurring pro-oncogenic mutations in EGFR and VEGFR2 (Figure S1e). However, we did not detect any high-frequency mutations that could spontaneously activate VEGFR1. We repeat our analysis using the KIRC patient data deposited in the cBioPortal repository^58^ (Figure 2d). In KIRC, all three RTKs exhibit very low mutation frequencies. Thus, our data analysis effectively rules out the possibility of a spontaneous VEGFR1 driver mutation in KIRC patients.

### VEGF_165_ induces transient phosphorylation of VEGFR1 in the plasma membrane and early endosome

Ligand-independent activation of RTKs in cancers overexpressing these receptors is a major driver of tumorigenesis^16,17,59^. Overexpression of the RTK induces concentration-dependent receptor dimerization at the plasma membrane, thereby enabling spontaneous autophosphorylation^14^. In contrast, VEGFR1 cannot be activated ligand-independently on the plasma membrane even when overexpressed, in our single-cell studies (Figures 3a-b and e)^9^. Profound JM inhibition prevents VEGFR1 ligand-independent dimerization, thereby stabilizing the monomeric conformation of the autoinhibited receptor^9^. It is unclear how the receptor density would modulate the ligand-dependent phosphorylation status of VEGFR1.

**Figure 3:**
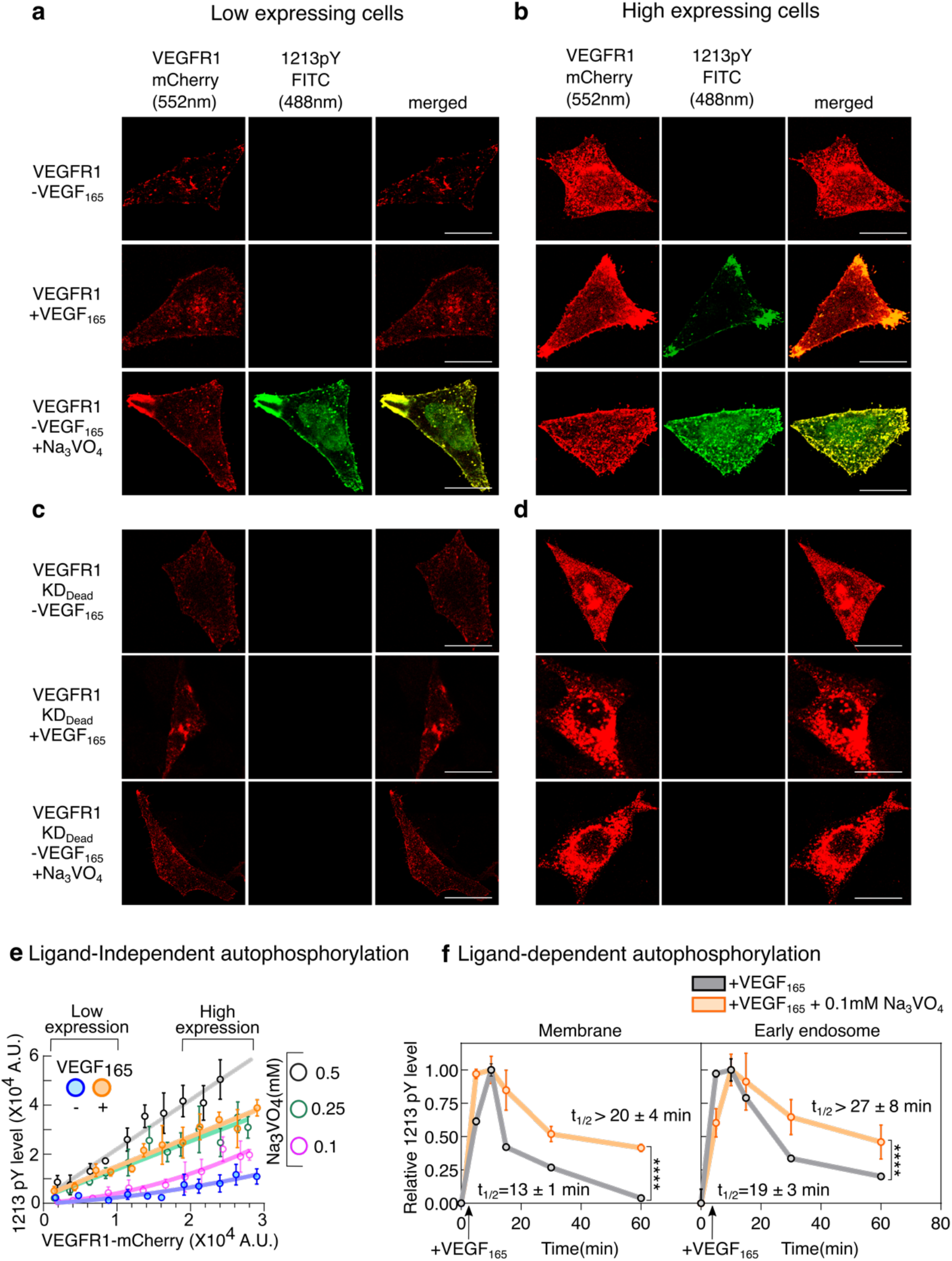
Measurement of the effect of pan-phosphatase inhibition on VEGFR1 ligand-independent or dependent autophosphorylation. **a-d)** Representative confocal images of the VEGFR1-mCherry constructs transiently expressed in CHO cells at the indicated expression levels. In each panel, cells were either activated with 100ng/ml of VEGF_165_ or with 0.1mM Na₃VO₄. Panels a and b represent wild-type VEGFR1, and panels c and d represent the kinase-dead D1022N mutant of VEGFR1. The red channel (λ_ex_ =552nm, λ_em_ = 570-660nm) denotes VEGFR1 expression. The green channel (λ_ex_ = 488nm, λ_em_ = 500-550nm) denotes Y1213 phosphorylation level (1213pY). The yellow represents the merged images. The cells with mCherry intensity between 0.1 × 10^4^ - 1 × 10^4^ AU at the plasma membrane were defined as low-expressing cells. Cells with mCherry intensity between 2 × 10^4^ to 3 × 10^4^ AU at the plasma membrane were defined as high-expressing cells. Scale bar = 10 µm. **e)** The VEGFR1-mCherry intensity at the plasma membrane is plotted against FITC intensity, denoting 1213pY level, n=120-150 cells. At each data point, cells were binned based on mCherry intensity. Each data point represents the mean ± SD from approximately 10–20 cells. The solid lines are guiding lines. The VEGFR1 was activated either with 100ng/ml of VEGF_165_ or at the indicated concentration of Na₃VO₄. **f)** The relative phosphorylation level of 1213Y (1213pY) measured at the plasma membrane (left) or at the early-endosome is plotted against time. The CHO cell line transiently expressing VEGFR1 was activated with 100 ng/ml VEGF_165_ (gray line) or 100 ng/ml VEGF_165_ in the presence of 0.1mM Na₃VO_4_ (orange line). Relative phosphorylation levels were determined by normalizing the background-corrected FITC: mCherry intensity ratio against the maximum. Each data point represents Mean ±SD from n=50-70 cells (n_total_ = 450-475 cells). Comparison between the two groups at the 60-minute time point was made using Student’s t-test. Statistical significance was assessed using the following criteria: \**p*< 0.05, \*\**p*< 0.01, \*\*\**p*< 0.001, \*\*\*\**p*< 0.0001, ns denotes not significant. **e-f)** The plots are made using Graphpad prism 8.0.2.

VEGFR1 binds VEGF-A with a ten-fold stronger affinity compared to VEGFR2^6–8^. The membrane-localized VEGFR1 in endothelial cells is long-lived (t_1/2_> 28 Hrs)^60^ compared to VEGFR2^61^. Therefore, we speculate that a long-lived VEGFR1 on the plasma membrane at higher receptor density would display a sustained phosphorylation upon ligand binding. To test, we turned to a single-cell assay to measure VEGFR1 phosphorylation at the plasma membrane and early endosomes (Figures 3f and S2h). We overexpressed VEGFR1-mCherry in the CHO (Chinese Hamster Ovary) cell line and measured the Y1213 phosphorylation (1213pY) at the C-terminal tail for sixty minutes after stimulating the cells with VEGF_165_. Using a single-cell assay, we measured 1213pY levels at the plasma membrane and in the early endosomes. The plasma membrane and the early endosome were labeled with WGA-Alexa Fluor 633 (blue), VEGFR1-mCherry expression indicated by red, and the phosphotyrosine 1213 (1213pY) is labeled green. As expected, we observed no change in VEGFR1 levels on the plasma membrane for one hour after activation (Figure S2g)^60^. Whereas, Y1213 at the C-terminal tail of VEGR1 was transiently phosphorylated with a half-life of 13 ± 1 min (Figure 3f and S2h). We observed a similar trend of transient Y1213 phosphorylation of VEGFR1 in the early endosome (t_1/2_= 19 ± 3 min). Very slow turnover of VEGFR1 at the plasma membrane (Figure S2g) suggests that the transient nature of tyrosine phosphorylation may reflect dephosphorylation rather than receptor degradation.

### Phosphatase inhibition remodels ligand-independent and ligand-dependent activation of VEGFR1

The phosphatases dampen downstream RTK signaling by removing phosphate groups from tyrosine residues^62^. Loss of phosphatase activity is often linked to activation of tyrosine kinases in multiple cancers^63,64^. Oxidative stress from reactive oxygen species (ROS) in the tumor microenvironment nonspecifically inhibits PTP, thereby activating VEGFR and other RTKs^65,66^. We ask whether inhibiting PTP with a nonspecific inhibitor, such as sodium orthovanadate (Na_3_VO_4_)^67^, is sufficient to remodel VEGFR1 phosphorylation.

We begin by measuring the ligand-independent phosphorylation of Y1213 in transiently transfected CHO cell lines expressing wild-type or kinase-dead mutant (D1022N) of VEGFR1-mCherry, respectively, after treatment with Na_3_VO_4_ (Figures 3a-e and S2a-f). We measure, in a single cell, the levels of plasma membrane-localized VEGFR1 from the mCherry intensity and extent of 1213pY from the green intensity. We observed that VEGFR1 is spontaneously autophosphorylated in the presence of 0.1 mM Na_3_VO_4_ and displays ligand-independent Y1213 phosphorylation (Figures 3a-b, e, and S2a). In a narrow concentration range of Na_3_VO_4_ (0.1 to 0.25mM), the Y1213 phosphorylation transitions to the linear pattern, as observed on ligand stimulation (Figures 3e and S2a). The kinase-dead mutant shows negligible Y1213 phosphorylation after PTP inhibition (Figures 3c-d and S3b-d). This independently confirms that inhibiting PTP increases the autophosphorylation of tyrosine residues in VEGFR1, and that this does not require another tyrosine kinase.

We next determine if inhibiting PTP in VEGFR1-overexpressing cells would remodel the time-dependent Y1213 phosphorylation. We measured the 1213pY levels in CHO cells overexpressing VEGFR1 (mCherry intensity of 1 to 2 × 10^4^ A.U.) over time (Figures 3f and S2e-h). The CHO cells transiently expressing VEGFR1 were activated with VEGF_165_ in the presence of 0.1mM Na_3_VO_4_. We observed that the rate of Y1213 phosphorylation at the plasma membrane and early endosomes remains unperturbed in cells treated with Na_3_VO_4_, compared to the control. However, the subtle inhibition of PTP (by 0.1 mM Na_3_VO_4_) increases the half-life (t_1/2_ > 20.0 ± 4 min at the plasma membrane and t_1/2_ > 27.0 ± 8 at the early-endosome) and amplitude of 1213pY levels (Figures 3f and S2e-f, h). We observed no change in the normalized mCherry intensity at the plasma membrane (Figure S2g), suggesting that VEGFR1 levels remain unchanged following Na_3_VO_4_ treatment. Our data indicate that inhibiting PTPs with Na_3_VO_4_ treatment is sufficient to remodel the VEGFR1 ligand-dependent and independent tyrosine phosphorylation. It is rather straightforward to explain how PTP inhibition remodels ligand-dependent VEGFR1 phosphorylation. But how does phosphatase inhibition activate the VEGFR1 ligand independently?

### Phosphatase inhibition induces spontaneous receptor dimerization

Receptor dimerization or oligomerization is a prerequisite for RTK activation^17,18,68^. The receptor dimerizes either at high receptor density at the plasma membrane or upon binding to its cognate ligands^14,30,69^. For VEGFR1, a stubborn JM inhibition prevents concentration-dependent receptor dimerization^9,56^. We next focus on understanding how a subtle change in the PTP activation profile perturbs the dynamics of VEGFR1 dimerization.

To measure VEGFR1 oligomerization, we turn to live-cell imaging of CHO cells transiently expressing VEGFR1 constructs tagged to mCherry by using two-dimensional stimulated emission depletion (2D STED) super-resolution microscopy. The VEGFR1 oligomers were determined from single-particle detection and tracking analysis. The oligomeric state was analyzed from the intensity distribution of VEGFR1 clusters (Figure 4b), the Diffusion coefficient derived from the mean-squared displacement (MSD) (Figures 4c-d, and S4), and step-wise photobleaching^70^ (Figure S5). In our experiment, we used two chimeric VEGFR1 constructs, VEGFR1-GPA and VEGFR1-G83I, as dimer and monomer controls, respectively (Figure 4a)^71^. The particle intensities of the monomer and dimer controls were used to optimize the linear density range to 0.15-0.25 particles/µm² (Figure S3). Within this particle density, the monomer control showed negligible spontaneous dimerization. The particle intensities for the monomer and dimer controls were fitted to single Gaussian components with mean intensities of 2.67 ± 1.89 A.U and 6.14 ± 2.55 A.U, respectively (Figure 4b and Table S6). The monomer and dimer controls were further validated by one-step and two-step photobleaching of the VEGFR1 construct, respectively (Figures S5a-b). The diffusion coefficient calculated from the MSD suggests that the monomer has a faster diffusion rate (0.487 ± 0.095 µm²s^-1^) compared to the dimer control (0.286 ± 0.087 µm²s^-1^) (Figures 4c-d). The single Gaussian fit for the ligand-free VEGFR1 clusters intensity, along with faster diffusion (D = 0.462 ± 0.117 µm²s^-1^) and one-step photobleaching, suggests that VEGFR1 is a monomer in the unligated state (Figures 4b-d and S5c). VEGF_165_ binding induces VEGFR1 dimerization, whose mixed Gaussian fitting yields two distinct components corresponding to monomer and dimer (Figure 4b). Ligand-dependent dimerization of VEGFR1 slows the diffusion rate (D = 0.354 ± 0.065 µm²s^-1^) (Figure 4c) and causes two-step photobleaching of the VEGFR1 cluster (Figure S5d). Unexpectedly, subtle inhibition of PTP with 0.1 mM Na_3_VO_4_ induces significant ligand-independent VEGFR1 dimerization (Figures 4b-d and S5e-f). Treating CHO cells with 0.1 mM Na_3_VO_4_ significantly reduces the diffusion speed of VEGFR1 clusters (D = 0.289 ± 0.112 µm²s^-1^) (Figures 4c-d). The presence of two components during mixed Gaussian fitting of particle intensities (Figure 4b) and two-step bleaching of the VEGFR1 cluster (Figures S5e-f) independently supports receptor dimerization. We conclude that the spontaneous ligand-independent dimerization of VEGFR1 upon PTP inhibition explains why VEGFR1 autophosphorylates at higher receptor density. We asked whether there is a specific PTP that is deregulated in KIRC?

**Figure 4:**
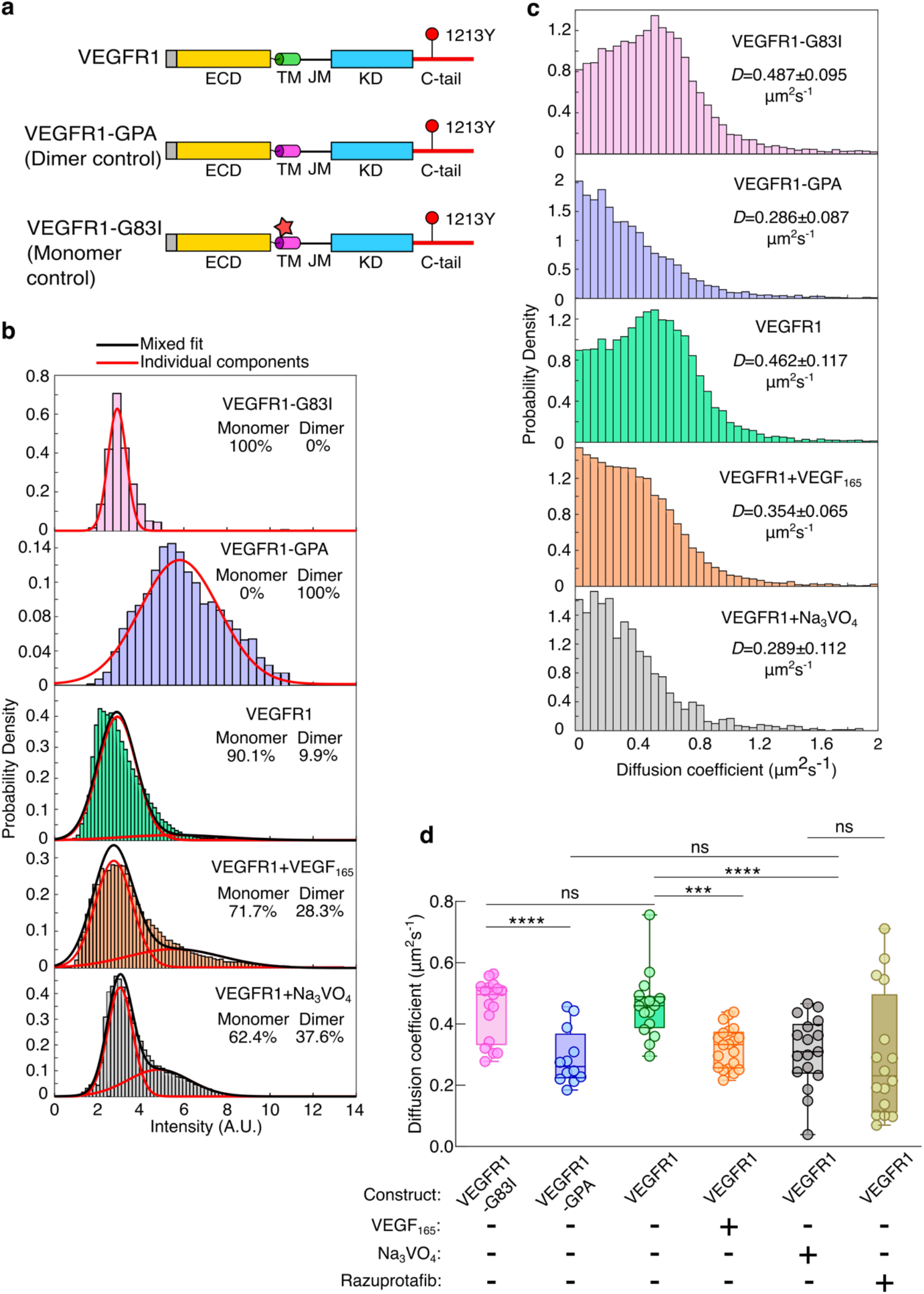
Probing dimerization of VEGFR1 on the plasma membrane. **a)** Schematic representation of VEGFR1 constructs used in this study. **b)** Histogram plot of intensity distributions for indicated VEGFR1 constructs against probability density. CHO cells transiently expressing VEGFR1 were activated with 100 ng/mL VEGF_165_, or 0.1 mM Na_3_VO_4_. Each panel represents the intensity measured for all particles in n=12-20 cells. The red and black lines show single-and mixed-Gaussian fits, respectively. The percentages of monomer and dimer populations for each panel are reported. **c)** Histogram distribution of diffusion coefficient for the indicated VEGFR1 constructs is plotted against probability density. Each panel represents the diffusion coefficient measured for all valid tracks in n=12-20 cells. The diffusion coefficient for each condition represents the median ± SD. **d)** The average diffusion coefficients of VEGFR1-mCherry measured from individual cells. Error bars represent mean ± SD from n = 12–20 cells. Comparison between the two groups was made using Student’s t-test. Statistical significance was assessed using the following criteria: \**p*< 0.05, \*\**p*< 0.01, \*\*\**p*< 0.001, \*\*\*\**p*< 0.0001, and ns denotes not significant. **a)** The schematic is made using Inkscape Ver1.4.3. **b-c)** The plot is made using MATLAB 2026a. **d)** The plot is made using Graphpad Prism 8.0.2.

### Gene Regulatory Networks reveal PTPRB phosphatase as a key regulator of VEGFR1 Phosphorylation

In the absence of the VEGFR1-phosphatase interactome, it is difficult to identify the potential PTP that dephosphorylates VEGFR1. Hence, we utilized a widely used computational method named “GENIE3” to infer two gene regulatory networks (GRNs), one from the transcriptomic profiles of the tumor tissue and another from those of the adjacent normal tissue of the KIRC patients^72^. Each GRN is a directed network in which a directed edge from gene A to gene B indicates that A (the regulator) regulates B (the target). For each edge, GENIE3 assigns a confidence score between 0 and 1, which represents GENIE3’s confidence in that edge. For each GRN, we extracted the subnetwork comprising the top 50 edges (by their confidence scores) where VEGFR1 is either the regulator or the target gene (Figures 5a-b). We noted that the PTPRB (also known as Vascular Endothelial Protein Tyrosine Phosphatase) is the only PTP that appeared to be a strong regulator of VEGFR1 function in tumor and adjacent normal tissue. To investigate further, we extracted the subnetworks comprising the top 50 edges in which VEGFR1 is either the regulator or the target of a receptor tyrosine phosphatase (Figures 5c-d). We noted that the PTPRB emerged as the highest-confidence phosphatase directly regulating VEGFR1. The loss of PTP often leads to activation of proto-oncogenic tyrosine kinases associated with malignant transformation in breast cancers and poor survival^63,73^. We wonder whether there is a correlation between PTPRB expression and disease prognosis in KIRC patients overexpressing VEGFR1.

**Figure 5:**
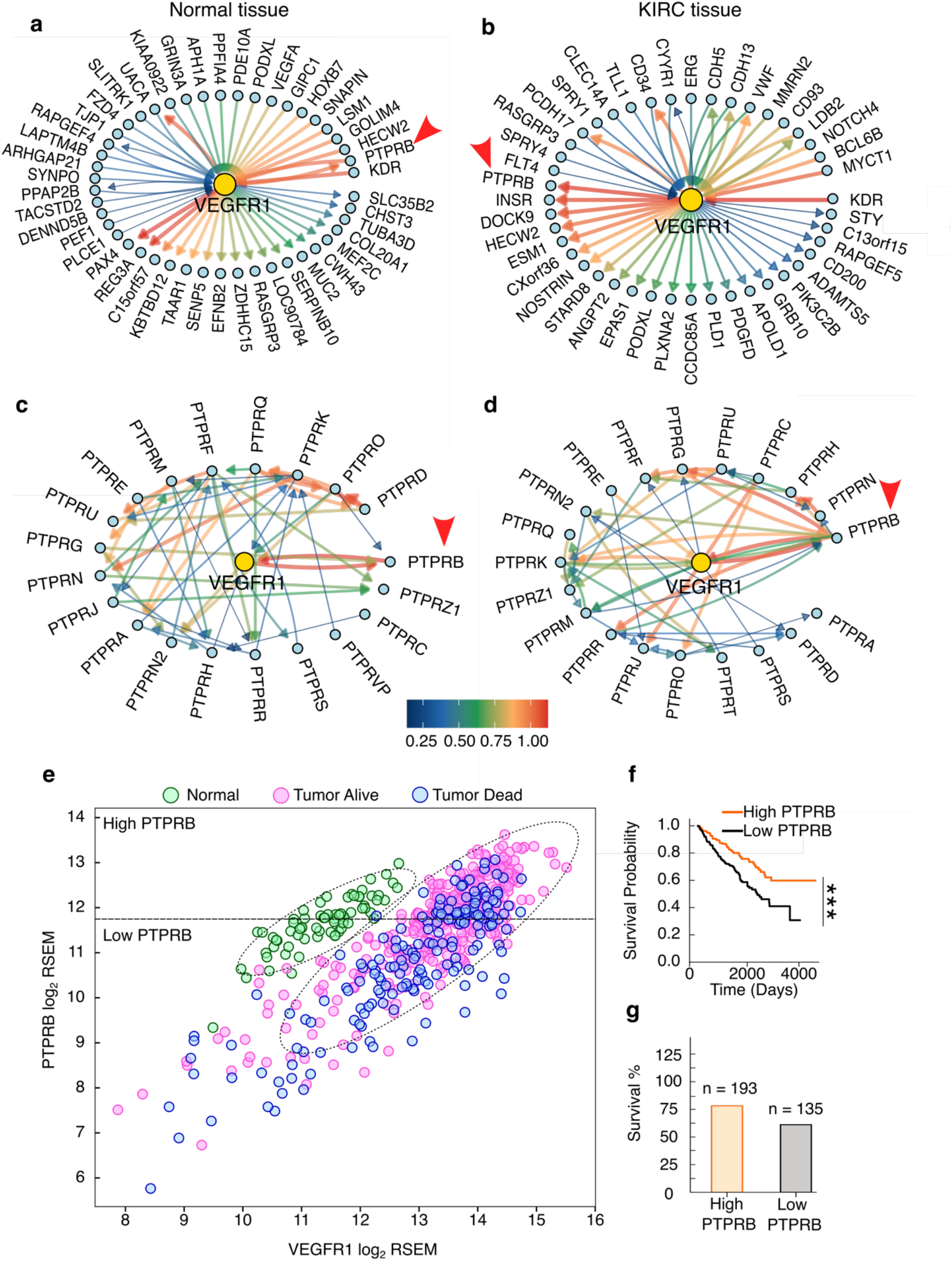
Gene regulatory network analysis of VEGFR1 in KIRC patients. **a-b)** Representative network highlighting the top 50 regulations involving VEGFR1 in normal and tumor tissue in KIRC patients, respectively. The direction of the arrow indicates the target gene. The color and width of the arrows indicate the relative strength of the regulation. **c-d)** Representative top 50 network highlighting the regulations between VEGFR1 and receptor tyrosine phosphatases in Normal tissue and tumor tissue in KIRC patients, respectively. **e)** A 2D correlation plot between PTPRB versus VEGFR1 expression levels (log_2_RSEM) in normal (green) and tumor (pink/blue) samples from KIRC patients. Each point denotes an individual patient sample. Based on TCGA sample codes and curated survival data, the tumor samples were classified into two categories: tumor alive (pink) (patients with higher survival), and tumor dead (blue) (patients with low survival). The area within the dotted lines denotes ellipses of maximum probability, representing 95% of the total patient population. The vertical dashed line represents median PTPRB expression. **f)** Kaplan-Meier plot comparing the probability of survival in patients with higher levels of PTPRB expression (orange line) versus lower levels of PTPRB expression (black line). A comparison between two groups was performed using Chi-square (*χ*^2^) test. Statistical significance was assessed using the following criteria: \**p*< 0.05, \*\**p*< 0.01, \*\*\**p*< 0.001, \*\*\*\**p*< 0.0001, and ns denotes not significant. **g)** Bar plot comparing the survival rate in patients with higher (orange) and lower PTPRB expression (grey). The survival percentage was calculated for each quadrant using the patient data within the ellipses of maximum probability. n denotes the number of patients survived, out of 247 and 221total patients in the high and low PTPRB groups, respectively. **a-g)** The plots are made using R 4.6.0.

### Loss of PTPRB expression in KIRC results in a poor survival rate

To determine a correlation between PTPRB and VEGFR1 expression to disease outcome, we plotted VEGFR1 mRNA expression against PTPRB transcription levels in tumor and adjacent normal tissue from KIRC patients. We observed that the tumor and the normal samples clustered distinctly (Figure 5e). Compared to the normal tissue samples, the correlation plot of VEGFR1 and PTPRB in the tumor tissue sample shows significantly greater dispersion. We estimate the survival probability from a Kaplan-Meier curve for KIRC patients expressing low or high PTPRB mRNA (Figure 5f). Our analysis indicates that patients overexpressing VEGFR1 along with low PTPRB levels have a lower survival rate (∼60%) than those with higher PTPRB levels (∼75%) (Figures 5f-g). Mapping the patients with low survival rates (blue circle) onto the VEGFR1-PTPRB correlation plot of tumor samples in Figure 5e clearly showed a poor disease prognosis for patients with less PTPRB expression. In conclusion, our analyses indicate that PTPRB expression levels may play a significant role in regulating VEGFR1 activation in tumors.

### Inhibition of PTPRB remodels VEGFR1 autophosphorylation

To verify whether inhibiting PTPRB activates VEGFR1, we recreated the single-cell assay system in the HEK293T cells. We chose HEK293T cells due to their low endogenous PTPRB expression (Figure S6e) and the absence of endogenous VEGFR1 or VEGFR2 expression. In our studies, we use Razuprotafib (also known as AKB-9778), a highly potent PTPRB inhibitor (IC_50_ = 17 pM)^74^. Using the single-cell assay, we first test the effect of PTPRB inhibition on ligand-independent VEGFR1 activation at the plasma membrane. Inhibiting PTPRB with Razuprotafib spontaneously activates the VEGFR1 at the plasma membrane (Figures 6 a-c). Between a narrow concentration range (0.0005-0.002 mM), Razuprotafib induces a receptor density-dependent VEGFR1 activation (Figure 6c), generally seen for EGFR and VEGFR2^9,14^. At a slightly higher Razuprotafib (0.005 mM) concentration, the VEGFR1 is linearly activated, similar to constitutively activating VEGFR2 mutant C482R linked to infantile hemangioma^9,75^. Our high-resolution image analysis of VEGFR1 shows that the Razuprotafib treatment (0.002 mM) induces robust ligand-independent receptor dimerization (Figures 6e-f and S6f-h). We observed that Razuprotafib treatment slows the diffusion rate of the VEGFR1 cluster (D = 0.287 ± 0.198 µm²s^-1^) (Figure 6f), similar to VEGFR1 dimer control (D = 0.286 ± 0.087 µm²s^-1^) (Figure 4c). The mixed Gaussian fitting of the VEGFR1 cluster intensity (Figure 6e) and two-step bleaching of the VEGFR1 cluster (Figure S6f) further support that inhibiting PTPRB induces spontaneous ligand-independent dimerization of VEGFR1.

**Figure 6:**
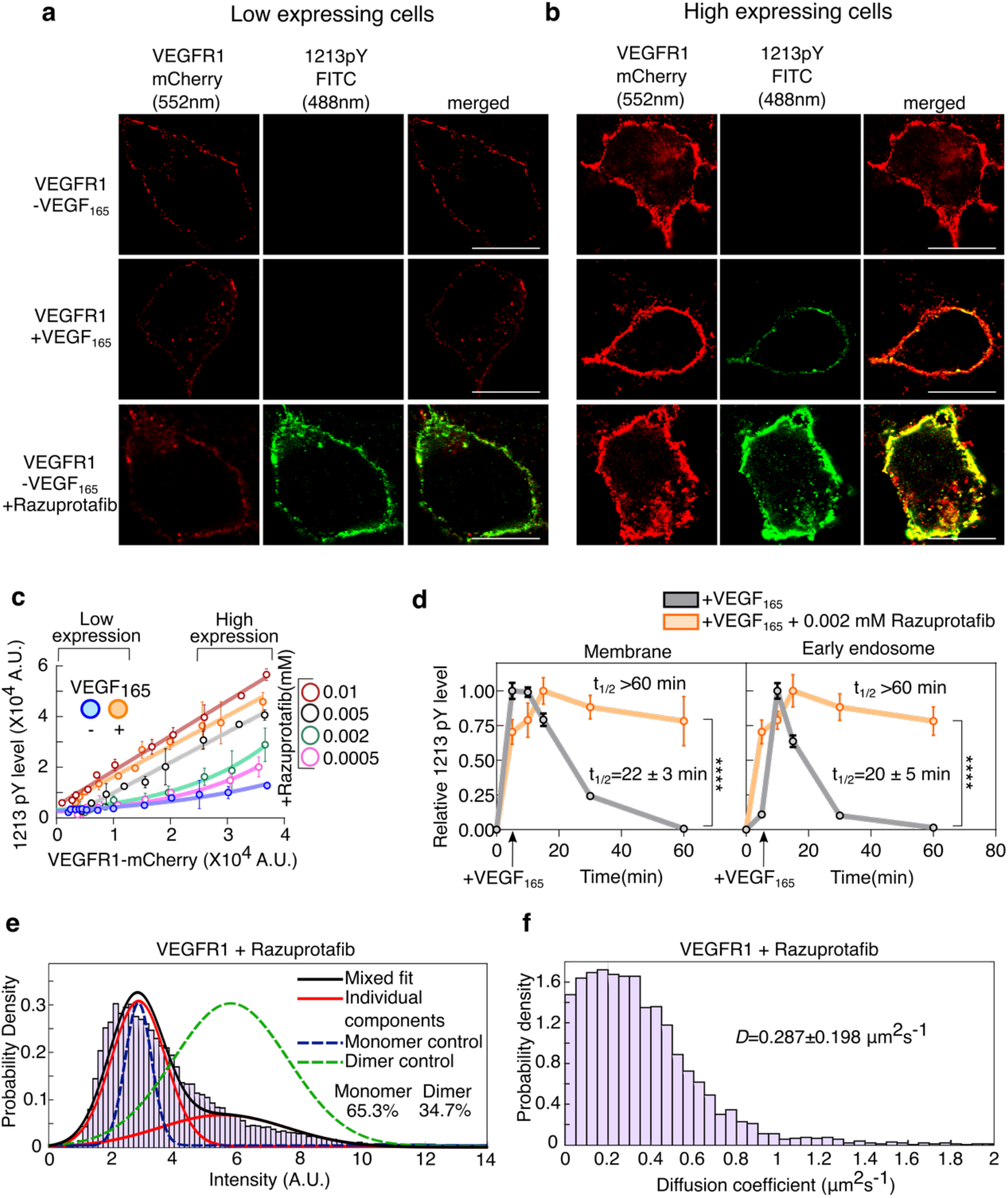
Effect of PTPRB inhibitor, Razuprotafib, on VEGFR1 oligomerization and autophosphorylation. **a-b)** Representative confocal images of HEK293T cells transiently expressing low (panel **a)** or high (panel **b)** levels of VEGFR1-mCherry. In each panel, cells were either activated with 100ng/ml of VEGF_165_ or with 0.002mM Razuprotafib. The red channel (λ_ex_ =552nm, λ_em_ = 570-660nm) denotes VEGFR1 expression. The green channel (λ_ex_ = 488nm, λ_em_ = 500-550nm) denotes Y1213 phosphorylation level (1213pY). The yellow represents the merged images. The cells with mCherry intensity between 0.1 × 10^4^ to 1 × 10^4^ AU at the plasma membrane were defined as low-expressing cells. Cells with mCherry intensity between 2.5 × 10^4^ to 4 × 10^4^ AU at the plasma membrane were defined as high-expressing cells. Scale bar = 10 µm. **c)** Intensity VEGFR1-mCherry at the plasma membrane of HEK293T is plotted against FITC intensity, denoting 1213pY level, n=130–180 cells. At each data point, cells were binned based on mCherry intensity. Each data point represents the mean ± SD from approximately 15–50 cells. The solid lines are guiding lines. The VEGFR1 was activated either with 100ng/ml of VEGF_165_ or at the indicated concentration of Razuprotafib. **d)** The relative phosphorylation level of 1213Y (1213pY) measured at the plasma membrane (left) or at the early-endosome is plotted against time. The HEK293T cell line transiently expressing VEGFR1 was activated with 100 ng/ml VEGF_165_ (gray line) or 100 ng/ml VEGF_165_ in the presence of 0.002mM Razuprotafib (orange line). Each data point represents. Mean ±SD from n=50-70 cells (n_total_ = 500-560 cells). A comparison between the two groups at the 60-minute time point was performed using Student’s t-test. Statistical significance was assessed using the following criteria: \**p*< 0.05, \*\**p*< 0.01, \*\*\**p*< 0.001, \*\*\*\**p*< 0.0001, and ns denotes not significant. **e)** Histogram distribution of diffusion coefficient for the VEGFR1 treated with 0.002mM Razuprotafib is plotted against probability density. The diffusion coefficient was measured for all valid tracks in HEK293T cells (n=12-20 cells). The diffusion coefficient represents the median ± SD. **f)** The intensity distributions for VEGFR1 treated with 0.002mM Razuprotafib is plotted against probability density. The histogram represents the intensity measured for all particles in n=12-20 cells. The blue and green lines show single-Gaussian fits for the monomer and dimer controls, respectively. The red and black lines show single-and mixed-Gaussian fits for VEGFR1 treated with 0.002mM Razuprotafib, respectively. The percentages of monomer and dimer populations are reported. **c-d)** The plots are made using Graphpad Prism 8.0.2. **e-f)** The plots are made using MATLAB 2026a.

As anticipated, Razuprotafib treatment remodels the ligand-dependent tyrosine phosphorylation of VEGFR1. Inhibiting PTPRB with Razuprotafib (0.002 mM) resulted in sustained tyrosine phosphorylation (t_1/2_ > 60 min) after VEGF_165_ stimulation, compared to transient phosphorylation of VEGFR1 observed in the absence of Razuprotafib treatment (Figures 6d and S6a-c). We observed that PTPRB inhibition does not alter the half-life (t_1/2_) of the VEGFR1 at the plasma membrane (Figure S6d). The change in phosphorylation lifetime may, in turn, determine cell fate^76^. Together, our data suggest that the PTPRB is a potential regulator of VEGFR1 activation and function.

### VEGFR2 ligand-dependent autophosphorylation is insensitive to PTPRB inhibition

PTPRB negatively regulates vascularization^77,78^ by dephosphorylating angioprotein receptor TIE2, VE-Cadherin, and VEGFR2^79–81^. Deletion or knockdown of PTPRB induces angiogenesis-like sprouting in human umbilical vein endothelial cells (HUVECs) or in mouse embryoid bodies^64,82^. Our multi-omics data analysis of TCGA and CPTAC data revealed that VEGFR2 is also overexpressed and robustly phosphorylated in KIRC (Figure S1). We previously showed that the ligand binding induced a sustained VEGFR2 autophosphorylation^9^. The receptor also has a higher propensity for ligand-dependent dimerization^9,18^. Thus, at higher receptor density, the VEGFR2 shows a spontaneous autophosphorylation. Our single-cell assay with transiently expressed VEGFR2 in HEK293T cells shows sustained Y1175 phosphorylation at the plasma membrane and early endosomes after ligand stimulation (Figures S7a-b). Inhibiting PTPRB with Razuprotafib increases the amplitude of Y1175 phosphorylation (Figures S7c-d), but does not change the half-life (t_1/2_) of phospho-tyrosine 1175 (Figures S7a-b). Our data suggest that at high receptor density, PTPRB does not influence the ligand-dependent activation of VEGFR2. However, a robust ligand-independent dimerization of VEGFR2 after phosphatase inhibition (Figure S8) indicates that the PTPRB inhibition may favor the ligand-independent activation of VEGFR2. In KIRC, transcriptomic analysis shows an overexpression of VEGF-A (Figure S1b). Therefore, we anticipate that in KIRC, PTPRB majorly remodels the ligand-dependent activation of VEGFR1.

## Discussion

KIRC is the most frequently diagnosed of all subtypes of kidney cancer, accounting for the majority of the cancer-related deaths^83^. The tumor is highly vascularized and shows overexpression of proangiogenic VEGFR1 and its ligand^43,44,84–86^. Knocking down VEGFR1 in the KIRC cell line results in a significant reduction in vascularization in the experimental mice xenograft model^26^. Several independent studies also support the importance of VEGFR1 in vascularization in multiple tumors^27,28^. Counterintuitively, during embryonic development, VEGFR1 negatively regulates angiogenesis by concealing VEGF-A from binding to the main angiogenic receptor VEGFR2^5,6,87^. Ligand binding induces a transient and weak tyrosine phosphorylation in VEGFR1^8,9^. The inhibitory JM-in conformation^37^ is stabilized by two main interactions: one is the docking of Y794 at the catalytic site as a pseudo-substrate. Second is the electrostatic interaction between the JM-S segment and the C-lobe of the kinase domain^9^. In the autoinhibited state, the Y794 may form a hydrogen bond with E878 in the C-helix^9,54^, which prevents the C-helix from swinging in and forming a conserved salt-bridge between K861 and E878^69,88^. The autoinhibited JM-in conformation prevents spontaneous receptor dimerization and activation at higher receptor density (Figures 3 and 4)^9^. Removing JM inhibition requires phosphorylation of Y794^89^ and unlocking of the electrostatic latch via ligand-dependent conformational rearrangement to the JM-out state^37^, allowing the receptor to dimerize and autophosphorylate (Figures 3 and 4).

The ligand-independent and dependent activation of VEGFR1 at higher receptor density may be explained by the equilibrium shift model involving multiple species (Figure 7)^9,14,30^. Here, we explain how strong JM inhibition (Figure 7)^9^ and PTPRB phosphatase activity (Figure 6) tame tyrosine phosphorylation of VEGFR1 under normal physiology (Figure 7)^9^. Downregulation of PTPRB activity due to Razuprotafib treatment (Figure 6) or loss of PTPRB expression (Figure 5e-g) is sufficient to release the JM-inhibition, leading to remodeling of ligand-independent and dependent phosphorylation of VEGFR1 (Figure 7). We speculate that the downregulation of PTPRB may stabilize phosphorylation at Y794, favoring receptor dimerization (Figures 4d and 6f) and activation (Figures 6a-c). VEGFR1 overexpression is emerging as a cancer cell marker that determines disease prognosis. The hyperactivation of VEGFR1 in KIRC patients overexpressing the receptor, and due to loss of PTPRB, may lead to increased tumor vascularization, which correlates with poor patient survival (Figures 5e-g). Our observation is consistent with the growing evidence suggesting that overexpression and activation of VEGFR1 are often associated with poor outcomes in cancer patients^90–92^.

**Figure 7:**
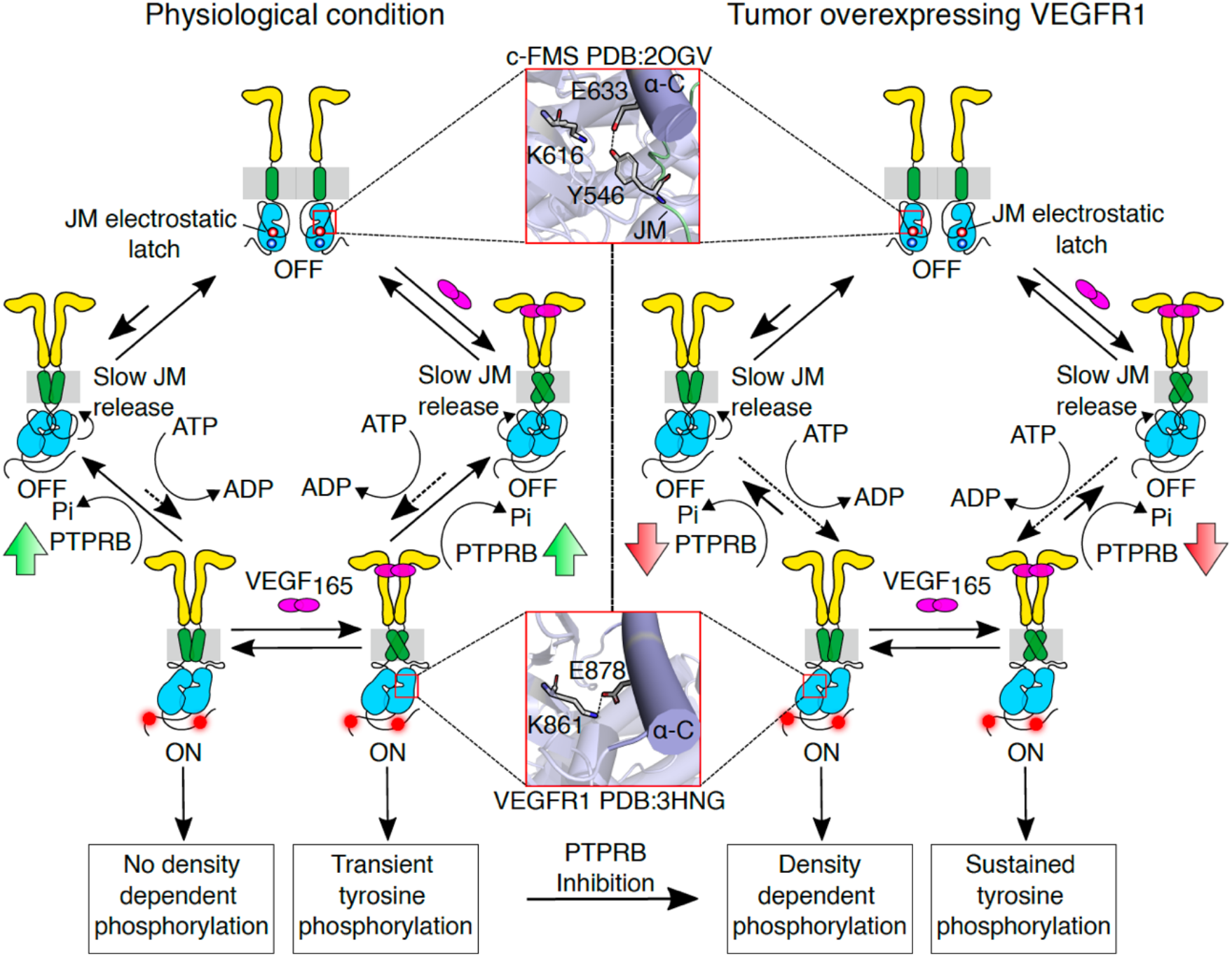
Proposed model for VEGFR1 regulation by phosphatase dynamics in KIRC. The left and right panels show the schematic representation of the VEGFR1 activation model in normal physiology and in tumors overexpressing the receptor. The various conformation species of VEGFR1 and corresponding activation states are labeled. The length of the black arrows indicates the receptor’s preferred state. The arrowhead connected to the dotted lines indicates the slow structural rearrangement from the inactive to the active state. Solid up (green) and down (red) arrows indicate high and low expression of PTPRB, respectively. The two inhibitory interactions, the JM electrostatic latch and the JM:αC interaction, which stabilizes the inactive state, are highlighted. The critical slat-bridge formed between the K861 and E878 in the αC in the active state is shown. The schematic is designed using Inkscape Ver1.4.3.

Tyrosine kinase signaling in metazoans has evolved as a write, read, and erase system^93^. While there are 58 receptor tyrosine kinases (writers)^30^, the signal is erased by 38 PTPs (erasers), of which only 21 are receptor-like PTPs^94^. The PTPs are promiscuous and can dephosphorylate a diverse set of tyrosine kinases^95^. Apart from VEGFR1, PTPRB regulates the phosphorylation status of multiple tyrosine kinases, like TIE2^74^, VEGFR2^82^, EGFR^62^, and FGFR^96^. Deregulation of PTPRB is thus linked to multiple malignant and non-malignant diseases^64,74,97–99^. On the other hand, the phosphorylation of one tyrosine kinase may be erased by multiple phosphatases. For example, phosphorylation of EGFR is regulated by PTPN12^63^, PTPRG/J and PTPN2^100^. The promiscuous relationship between RTKs and PTPs network^62^ may enable cells to reprogram phosphatase expression to prevent nonspecific activation of RTKs. This explains why, in CHO cells that do not express PTPRB, the VEGFR1 phosphorylation is downregulated.

The development of PTP inhibitors targeting a specific RTK signaling pathway faces multiple challenges^101^. Nevertheless, PTPRB inhibitors such as Razuprotafib have been evaluated clinically for the treatment of vascular disorders in diabetic retinopathy^74^ and glaucoma^96^. Additionally, restoring the tumor-suppressor activity of protein phosphatase 2A (PP2A) with a small-molecule activator showed promising results in treating various drug-resistant cancers^102,103^. In conclusion, we have presented a general mechanism that explains how loss of PTP activates the benign tyrosine kinase VEGFR1, thereby driving tumor metastasis. Phosphatase deficiency promotes VEGFR1 dimerization by favoring the transition to an active conformation. Developing PTP activators presents an opportunity to revert VEGFR1 to a stable inactive state.

## Experimental Procedures

The experimental procedure and material section are provided in the Supplemental Data.

## Supplemental Data

The Supplemental Data contains Experimental Procedures, Figure S1-S8, and Table S1-S6.

## Acknowledgments

The authors thank Prof. Bidisha Sinha and Prof. Arnab Gupta for access to the microscopy facility and for helpful discussions. The mutation data was obtained from the Sanger Institute Catalog Of Somatic Mutations In Cancer web site, http://cancer.sanger.ac.uk/cosmic.

## Author Contributions

The manuscript was written through the contribution of all authors. All authors have approved the final version of the manuscript. RD and SG designed the experiments. SG, MPC, and BD performed the Biochemical experiments, Imaging, and data analysis. AP performed the patient data analysis; SP supervised and guided GRN analysis. AG performed the image analysis. RD and SG wrote the manuscript.

## Funding and additional information

The authors thank IISER Kolkata for research funding, CIF, IISER Kolkata, for instrument facilities, and the DBT Builder Project (BT/INF/22/SP45383/2022). This work is supported by a grant from ANRF (ANRF/ARF/2025/000629/LS). Fellowships from CSIR-UGC support MPC and BD.

## Data Availability Statement

All the relevant data are contained within this article and in the supplemental data. Source data are provided with this paper in the Source Data file. Uncropped blots are available in the Source Data file. Source data are provided with this paper.

## References

1 Simons, M., Gordon, E. & Claesson-Welsh, L. Mechanisms and regulation of endothelial VEGF receptor signalling. Nat Rev Mol Cell Biol 17, 611–625, doi:10.1038/nrm.2016.87 (2016).

2 Millauer, B. et al. High affinity VEGF binding and developmental expression suggest Flk-1 as a major regulator of vasculogenesis and angiogenesis. Cell 72, 835–846, doi:10.1016/0092-8674(93)90573-9 (1993).

3 Shibuya, M. et al. Nucleotide sequence and expression of a novel human receptor-type tyrosine kinase gene (flt) closely related to the fms family. Oncogene 5, 519–524 (1990).

4 de Vries, C. et al. The fms-like tyrosine kinase, a receptor for vascular endothelial growth factor. Science 255, 989–991, doi:10.1126/science.1312256 (1992).

5 Park, J. E., Chen, H. H., Winer, J., Houck, K. A. & Ferrara, N. Placenta growth factor. Potentiation of vascular endothelial growth factor bioactivity, in vitro and in vivo, and high affinity binding to Flt-1 but not to Flk-1/KDR. J Biol Chem 269, 25646–25654 (1994).

6 Hiratsuka, S., Minowa, O., Kuno, J., Noda, T. & Shibuya, M. Flt-1 lacking the tyrosine kinase domain is sufficient for normal development and angiogenesis in mice. Proc Natl Acad Sci U S A 95, 9349–9354, doi:10.1073/pnas.95.16.9349 (1998).

7 Shinkai, A. et al. Mapping of the sites involved in ligand association and dissociation at the extracellular domain of the kinase insert domain-containing receptor for vascular endothelial growth factor. J Biol Chem 273, 31283–31288, doi:10.1074/jbc.273.47.31283 (1998).

8 Waltenberger, J., Claesson-Welsh, L., Siegbahn, A., Shibuya, M. & Heldin, C. H. Different signal transduction properties of KDR and Flt1, two receptors for vascular endothelial growth factor. Journal of Biological Chemistry 269, 26988–26995, doi:10.1016/s0021-9258(18)47116-5 (1994).

9 Chakraborty, M. P. et al. Molecular basis of VEGFR1 autoinhibition at the plasma membrane. Nat Commun 15, 1346, doi:10.1038/s41467-024-45499-2 (2024).

10 A Sawano 1, T. T., S Yamaguchi, M Aonuma, M Shibuya. Flt-1 but not KDR/Flk-1 tyrosine kinase is a receptor for placenta growth factor, which is related to vascular endothelial growth factor. Cell Growth Differ. 1996, PMID: 8822205

11 L, S., 1 &, G. N., Maru Y, Neufeld G, Yamaguchi S, Shibuya M. A unique signal transduction from FLT tyrosine kinase, a receptor for vascular endothelial growth factor VEGF. Oncogene. 1995, PMID: 7824266

12 Barleon, B. et al. Migration of human monocytes in response to vascular endothelial growth factor (VEGF) is mediated via the VEGF receptor flt-1. Blood 87, 3336–3343, doi:10.1182/blood.V87.8.3336.bloodjournal8783336 (1996).

13 Hirsch, F. R., Varella-Garcia, M. & Cappuzzo, F. Predictive value of EGFR and HER2 overexpression in advanced non-small-cell lung cancer. Oncogene 28 **Suppl 1**, S32–37, doi:10.1038/onc.2009.199 (2009).

14 Endres, N. F. et al. Conformational coupling across the plasma membrane in activation of the EGF receptor. Cell 152, 543–556, doi:10.1016/j.cell.2012.12.032 (2013).

15 Du, Z. & Lovly, C. M. Mechanisms of receptor tyrosine kinase activation in cancer. Mol Cancer 17, 58, doi:10.1186/s12943-018-0782-4 (2018).

16 Sarabipour, S. & Hristova, K. Mechanism of FGF receptor dimerization and activation. Nat Commun 7, 10262, doi:10.1038/ncomms10262 (2016).

17 Chung, I. et al. Spatial control of EGF receptor activation by reversible dimerization on living cells. Nature 464, 783–787, doi:10.1038/nature08827 (2010).

18 Sarabipour, S., Ballmer-Hofer, K. & Hristova, K. VEGFR-2 conformational switch in response to ligand binding. Elife 5, e13876, doi:10.7554/eLife.13876 (2016).

19 Hiratsuka, S. et al. Involvement of Flt-1 tyrosine kinase (vascular endothelial growth factor receptor-1) in pathological angiogenesis. Cancer Res 61, 1207–1213 (2001).

20 Bates, R. C. et al. Flt-1-dependent survival characterizes the epithelial-mesenchymal transition of colonic organoids. Curr Biol 13, 1721–1727, doi:10.1016/j.cub.2003.09.002 (2003).

21 Kaplan, R. N., Psaila, B. & Lyden, D. Bone marrow cells in the ’pre-metastatic niche’: within bone and beyond. Cancer Metastasis Rev 25, 521–529, doi:10.1007/s10555-006-9036-9 (2006).

22 Fischer, C., Mazzone, M., Jonckx, B. & Carmeliet, P. FLT1 and its ligands VEGFB and PlGF: drug targets for anti-angiogenic therapy? Nat Rev Cancer 8, 942–956, doi:10.1038/nrc2524 (2008).

23 Cao, R. et al. VEGFR1-mediated pericyte ablation links VEGF and PlGF to cancer-associated retinopathy. Proc Natl Acad Sci U S A 107, 856–861, doi:10.1073/pnas.0911661107 (2010).

24 Qian, B. Z. et al. CCL2 recruits inflammatory monocytes to facilitate breast-tumour metastasis. Nature 475, 222–225, doi:10.1038/nature10138 (2011).

25 Qian, B. Z. et al. FLT1 signaling in metastasis-associated macrophages activates an inflammatory signature that promotes breast cancer metastasis. J Exp Med 212, 1433–1448, doi:10.1084/jem.20141555 (2015).

26 Li, C., Liu, B., Dai, Z. & Tao, Y. Knockdown of VEGF receptor-1 (VEGFR-1) impairs macrophage infiltration, angiogenesis and growth of clear cell renal cell carcinoma (CRCC). Cancer Biol Ther 12, 872–880, doi:10.4161/cbt.12.10.17672 (2011).

27 Kerber, M. et al. Flt-1 signaling in macrophages promotes glioma growth in vivo. Cancer Res 68, 7342–7351, doi:10.1158/0008-5472.CAN-07-6241 (2008).

28 Fischer, C. et al. Anti-PlGF inhibits growth of VEGF(R)-inhibitor-resistant tumors without affecting healthy vessels. Cell 131, 463–475, doi:10.1016/j.cell.2007.08.038 (2007).

29 Selvaraj, D. et al. A Functional Role for VEGFR1 Expressed in Peripheral Sensory Neurons in Cancer Pain. Cancer Cell 27, 780–796, doi:10.1016/j.ccell.2015.04.017 (2015).

30 Lemmon, M. A. & Schlessinger, J. Cell signaling by receptor tyrosine kinases. Cell 141, 1117–1134, doi:10.1016/j.cell.2010.06.011 (2010).

31 Olsson, A. K., Dimberg, A., Kreuger, J. & Claesson-Welsh, L. VEGF receptor signalling - in control of vascular function. Nat Rev Mol Cell Biol 7, 359–371, doi:10.1038/nrm1911 (2006).

32 Yang, Y., Xie, P., Opatowsky, Y. & Schlessinger, J. Direct contacts between extracellular membrane-proximal domains are required for VEGF receptor activation and cell signaling. Proc Natl Acad Sci U S A 107, 1906–1911, doi:10.1073/pnas.0914052107 (2010).

33 Ahmadova, Z. et al. Fluorescent Resonance Energy Transfer Imaging of VEGFR Dimerization. Anticancer Research 34, 2123–2133 (2014).

34 Tao, Q., Backer, M. V., Backer, J. M. & Terman, B. I. Kinase insert domain receptor (KDR) extracellular immunoglobulin-like domains 4-7 contain structural features that block receptor dimerization and vascular endothelial growth factor-induced signaling. J Biol Chem 276, 21916–21923, doi:10.1074/jbc.M100763200 (2001).

35 Yuzawa, S. et al. Structural basis for activation of the receptor tyrosine kinase KIT by stem cell factor. Cell 130, 323–334, doi:10.1016/j.cell.2007.05.055 (2007).

36 Hubbard, S. R. Juxtamembrane autoinhibition in receptor tyrosine kinases. Nat Rev Mol Cell Biol 5, 464–471, doi:10.1038/nrm1399 (2004).

37 McTigue, M. et al. Molecular conformations, interactions, and properties associated with drug efficiency and clinical performance among VEGFR TK inhibitors. Proc Natl Acad Sci U S A 109, 18281–18289, doi:10.1073/pnas.1207759109 (2012).

38 Yarden, Y. & Schlessinger, J. Epidermal growth factor induces rapid, reversible aggregation of the purified epidermal growth factor receptor. Biochemistry 26, 1443–1451, doi:10.1021/bi00379a035 (1987).

39 Huang, Y. et al. Molecular basis for multimerization in the activation of the epidermal growth factor receptor. Elife 5, doi:10.7554/eLife.14107 (2016).

40 Markovic-Mueller, S. et al. Structure of the Full-length VEGFR-1 Extracellular Domain in Complex with VEGF-A. Structure 25, 341–352, doi:10.1016/j.str.2016.12.012 (2017).

41 Cancer Genome Atlas Research, N., et al. The Cancer Genome Atlas Pan-Cancer analysis project. Nat Genet 45, 1113–1120, doi:10.1038/ng.2764 (2013).

42 Cancer Genome Atlas Research, N. Comprehensive molecular characterization of clear cell renal cell carcinoma. Nature 499, 43–49, doi:10.1038/nature12222 (2013).

43 Clark, D. J. et al. Integrated Proteogenomic Characterization of Clear Cell Renal Cell Carcinoma. Cell 179, 964–983 e931, doi:10.1016/j.cell.2019.10.007 (2019).

44 Quail, D. F. & Joyce, J. A. Microenvironmental regulation of tumor progression and metastasis. Nat Med 19, 1423–1437, doi:10.1038/nm.3394 (2013).

45 Gerber, H. P., Condorelli, F., Park, J. & Ferrara, N. Differential transcriptional regulation of the two vascular endothelial growth factor receptor genes. Flt-1, but not Flk-1/KDR, is up-regulated by hypoxia. J Biol Chem 272, 23659–23667, doi:10.1074/jbc.272.38.23659 (1997).

46 van Houwelingen, K. P. et al. Prevalence of von Hippel-Lindau gene mutations in sporadic renal cell carcinoma: results from The Netherlands cohort study. BMC Cancer 5, 57, doi:10.1186/1471-2407-5-57 (2005).

47 Igarashi, H., Esumi, M., Ishida, H. & Okada, K. Vascular endothelial growth factor overexpression is correlated with von Hippel-Lindau tumor suppressor gene inactivation in patients with sporadic renal cell carcinoma. Cancer 95, 47–53, doi:10.1002/cncr.10635 (2002).

48 Rahimi, N., Dayanir, V. & Lashkari, K. Receptor chimeras indicate that the vascular endothelial growth factor receptor-1 (VEGFR-1) modulates mitogenic activity of VEGFR-2 in endothelial cells. J Biol Chem 275, 16986–16992, doi:10.1074/jbc.M000528200 (2000).

49 Shibuya, M. Vascular endothelial growth factor receptor-1 (VEGFR-1/Flt-1): a dual regulator for angiogenesis. Angiogenesis 9, 225–230; discussion 231, doi:10.1007/s10456-006-9055-8 (2006).

50 Vogelstein, B. et al. Cancer genome landscapes. Science 339, 1546–1558, doi:10.1126/science.1235122 (2013).

51 Silva, I. P. et al. Identification of a Novel Pathogenic Germline KDR Variant in Melanoma. Clin Cancer Res 22, 2377–2385, doi:10.1158/1078-0432.CCR-15-1811 (2016).

52 Binder, Z. A. et al. Epidermal Growth Factor Receptor Extracellular Domain Mutations in Glioblastoma Present Opportunities for Clinical Imaging and Therapeutic Development. Cancer Cell 34, 163–177 e167, doi:10.1016/j.ccell.2018.06.006 (2018).

53 Yin, L. X. et al. Prognostic impact and treatment options for EGFR-mutant non-small cell lung cancer with concurrent EGFR amplification. Lung Cancer 214, 109339, doi:10.1016/j.lungcan.2026.109339 (2026).

54 Walter, M. et al. The 2.7 A crystal structure of the autoinhibited human c-Fms kinase domain. J Mol Biol 367, 839–847, doi:10.1016/j.jmb.2007.01.036 (2007).

55 Meyer, R. D., Mohammadi, M. & Rahimi, N. A single amino acid substitution in the activation loop defines the decoy characteristic of VEGFR-1/FLT-1. J Biol Chem 281, 867–875, doi:10.1074/jbc.M506454200 (2006).

56 Gille, H. et al. A repressor sequence in the juxtamembrane domain of Flt-1 (VEGFR-1) constitutively inhibits vascular endothelial growth factor-dependent phosphatidylinositol 3’-kinase activation and endothelial cell migration. EMBO J 19, 4064–4073, doi:10.1093/emboj/19.15.4064 (2000).

57 Bamford, S. et al. The COSMIC (Catalogue of Somatic Mutations in Cancer) database and website. Br J Cancer 91, 355–358, doi:10.1038/sj.bjc.6601894 (2004).

58 Miao, D. et al. Genomic correlates of response to immune checkpoint therapies in clear cell renal cell carcinoma. Science 359, 801–806, doi:10.1126/science.aan5951 (2018).

59 Lin, C. C. et al. Inhibition of basal FGF receptor signaling by dimeric Grb2. Cell 149, 1514–1524, doi:10.1016/j.cell.2012.04.033 (2012).

60 Boucher, J. M. et al. Dynamic alterations in decoy VEGF receptor-1 stability regulate angiogenesis. Nat Commun 8, 15699, doi:10.1038/ncomms15699 (2017).

61 Yen, H. C., Xu, Q., Chou, D. M., Zhao, Z. & Elledge, S. J. Global protein stability profiling in mammalian cells. Science 322, 918–923, doi:10.1126/science.1160489 (2008).

62 Yao, Z. et al. A Global Analysis of the Receptor Tyrosine Kinase-Protein Phosphatase Interactome. Mol Cell 65, 347–360, doi:10.1016/j.molcel.2016.12.004 (2017).

63 Sun, T. et al. Activation of multiple proto-oncogenic tyrosine kinases in breast cancer via loss of the PTPN12 phosphatase. Cell 144, 703–718, doi:10.1016/j.cell.2011.02.003 (2011).

64 Ramo, J. T. et al. Rare genetic variation in PTPRB is associated with central serous chorioretinopathy, varicose veins and glaucoma. Nat Commun 16, 4127, doi:10.1038/s41467-025-58686-6 (2025).

65 Bae, Y. S. et al. Epidermal growth factor (EGF)-induced generation of hydrogen peroxide - Role in EGF receptor-mediated tyrosine phosphorylation. Journal of Biological Chemistry 272, 217–221, doi:DOI 10.1074/jbc.272.1.217 (1997).

66 Sundaresan, M., Yu, Z. X., Ferrans, V. J., Irani, K. & Finkel, T. Requirement for generation of H2O2 for platelet-derived growth factor signal transduction. Science 270, 296–299, doi:10.1126/science.270.5234.296 (1995).

67 Hecht, D. & Zick, Y. Selective inhibition of protein tyrosine phosphatase activities by H2O2 and vanadate in vitro. Biochem Biophys Res Commun 188, 773–779, doi:10.1016/0006-291x(92)91123-8 (1992).

68 Plotnikov, A. N., Schlessinger, J., Hubbard, S. R. & Mohammadi, M. Structural basis for FGF receptor dimerization and activation. Cell 98, 641–650, doi:10.1016/s0092-8674(00)80051-3 (1999).

69 Jura, N. et al. Catalytic control in the EGF receptor and its connection to general kinase regulatory mechanisms. Mol Cell 42, 9–22, doi:10.1016/j.molcel.2011.03.004 (2011).

70 Calebiro, D. et al. Single-molecule analysis of fluorescently labeled G-protein-coupled receptors reveals complexes with distinct dynamics and organization. Proc Natl Acad Sci U S A 110, 743–748, doi:10.1073/pnas.1205798110 (2013).

71 Lemmon, M. A., Flanagan, J. M., Treutlein, H. R., Zhang, J. & Engelman, D. M. Sequence specificity in the dimerization of transmembrane alpha-helices. Biochemistry 31, 12719–12725, doi:10.1021/bi00166a002 (1992).

72 Huynh-Thu, V. A., Irrthum, A., Wehenkel, L. & Geurts, P. Inferring regulatory networks from expression data using tree-based methods. PLoS One 5, doi:10.1371/journal.pone.0012776 (2010).

73 Dong, H. et al. PTPRO represses ERBB2-driven breast oncogenesis by dephosphorylation and endosomal internalization of ERBB2. Oncogene 36, 410–422, doi:10.1038/onc.2016.213 (2017).

74 Shen, J. et al. Targeting VE-PTP activates TIE2 and stabilizes the ocular vasculature. J Clin Invest 124, 4564–4576, doi:10.1172/JCI74527 (2014).

75 Jinnin, M. et al. Suppressed NFAT-dependent VEGFR1 expression and constitutive VEGFR2 signaling in infantile hemangioma. Nat Med 14, 1236–1246, doi:10.1038/nm.1877 (2008).

76 Kaplan, M. et al. EGFR Dynamics Change during Activation in Native Membranes as Revealed by NMR. Cell 167, 1241–1251 e1211, doi:10.1016/j.cell.2016.10.038 (2016).

77 Baumer, S. et al. Vascular endothelial cell-specific phosphotyrosine phosphatase (VE-PTP) activity is required for blood vessel development. Blood 107, 4754–4762, doi:10.1182/blood-2006-01-0141 (2006).

78 Dominguez, M. G. et al. Vascular endothelial tyrosine phosphatase (VE-PTP)-null mice undergo vasculogenesis but die embryonically because of defects in angiogenesis. Proc Natl Acad Sci U S A 104, 3243–3248, doi:10.1073/pnas.0611510104 (2007).

79 Fachinger, G., Deutsch, U. & Risau, W. Functional interaction of vascular endothelial-protein-tyrosine phosphatase with the angiopoietin receptor Tie-2. Oncogene 18, 5948–5953, doi:10.1038/sj.onc.1202992 (1999).

80 Mellberg, S. et al. Transcriptional profiling reveals a critical role for tyrosine phosphatase VE-PTP in regulation of VEGFR2 activity and endothelial cell morphogenesis. FASEB J 23, 1490–1502, doi:10.1096/fj.08-123810 (2009).

81 Nottebaum, A. F. et al. VE-PTP maintains the endothelial barrier via plakoglobin and becomes dissociated from VE-cadherin by leukocytes and by VEGF. J Exp Med 205, 2929–2945, doi:10.1084/jem.20080406 (2008).

82 Hayashi, M. et al. VE-PTP regulates VEGFR2 activity in stalk cells to establish endothelial cell polarity and lumen formation. Nat Commun 4, 1672, doi:10.1038/ncomms2683 (2013).

83 Hsieh, J. J. et al. Renal cell carcinoma. Nat Rev Dis Primers 3, 17009, doi:10.1038/nrdp.2017.9 (2017).

84 Rivet, J. et al. VEGF and VEGFR-1 are coexpressed by epithelial and stromal cells of renal cell carcinoma. Cancer 112, 433–442, doi:10.1002/cncr.23186 (2008).

85 Qu, Y. et al. Proteogenomic characterization of MiT family translocation renal cell carcinoma. Nat Commun 13, 7494, doi:10.1038/s41467-022-34460-w (2022).

86 Kaplan, R. N. et al. VEGFR1-positive haematopoietic bone marrow progenitors initiate the pre-metastatic niche. Nature 438, 820–827, doi:10.1038/nature04186 (2005).

87 Fong, G. H., Rossant, J., Gertsenstein, M. & Breitman, M. L. Role of the Flt-1 receptor tyrosine kinase in regulating the assembly of vascular endothelium. Nature 376, 66–70, doi:10.1038/376066a0 (1995).

88 Taylor, S. S. et al. From structure to the dynamic regulation of a molecular switch: A journey over 3 decades. J Biol Chem 296, 100746, doi:10.1016/j.jbc.2021.100746 (2021).

89 Saitoh, M. et al. Identification of important regions in the cytoplasmic juxtamembrane domain of type I receptor that separate signaling pathways of transforming growth factor-beta. J Biol Chem 271, 2769–2775, doi:10.1074/jbc.271.5.2769 (1996).

90 Yang, X. et al. VEGF-B promotes cancer metastasis through a VEGF-A-independent mechanism and serves as a marker of poor prognosis for cancer patients. Proc Natl Acad Sci U S A 112, E2900–2909, doi:10.1073/pnas.1503500112 (2015).

91 Freire Valls, A., et al. VEGFR1(+) Metastasis-Associated Macrophages Contribute to Metastatic Angiogenesis and Influence Colorectal Cancer Patient Outcome. Clin Cancer Res 25, 5674–5685, doi:10.1158/1078-0432.CCR-18-2123 (2019).

92 Fragoso, R. et al. VEGFR-1 (FLT-1) activation modulates acute lymphoblastic leukemia localization and survival within the bone marrow, determining the onset of extramedullary disease. Blood 107, 1608–1616, doi:10.1182/blood-2005-06-2530 (2006).

93 Lim, W. A. & Pawson, T. Phosphotyrosine signaling: evolving a new cellular communication system. Cell 142, 661–667, doi:10.1016/j.cell.2010.08.023 (2010).

94 Alonso, A. et al. Protein tyrosine phosphatases in the human genome. Cell 117, 699–711, doi:10.1016/j.cell.2004.05.018 (2004).

95 Pincus, D., Letunic, I., Bork, P. & Lim, W. A. Evolution of the phospho-tyrosine signaling machinery in premetazoan lineages. Proc Natl Acad Sci U S A 105, 9680–9684, doi:10.1073/pnas.0803161105 (2008).

96 Thomson, B. R. et al. Targeting the vascular-specific phosphatase PTPRB protects against retinal ganglion cell loss in a pre-clinical model of glaucoma. Elife 8, doi:10.7554/eLife.48474 (2019).

97 Behjati, S. et al. Recurrent PTPRB and PLCG1 mutations in angiosarcoma. Nat Genet 46, 376–379, doi:10.1038/ng.2921 (2014).

98 Weng, X. et al. PTPRB promotes metastasis of colorectal carcinoma via inducing epithelial-mesenchymal transition. Cell Death Dis 10, 352, doi:10.1038/s41419-019-1554-9 (2019).

99 Carota, I. A. et al. Targeting VE-PTP phosphatase protects the kidney from diabetic injury. J Exp Med 216, 936–949, doi:10.1084/jem.20180009 (2019).

100 Stanoev, A. et al. Interdependence between EGFR and Phosphatases Spatially Established by Vesicular Dynamics Generates a Growth Factor Sensing and Responding Network. Cell Syst 7, 295–309 e211, doi:10.1016/j.cels.2018.06.006 (2018).

101 Tonks, N. K. Protein tyrosine phosphatases: from genes, to function, to disease. Nat Rev Mol Cell Biol 7, 833–846, doi:10.1038/nrm2039 (2006).

102 Neviani, P. et al. FTY720, a new alternative for treating blast crisis chronic myelogenous leukemia and Philadelphia chromosome-positive acute lymphocytic leukemia. J Clin Invest 117, 2408–2421, doi:10.1172/JCI31095 (2007).

103 McClinch, K. et al. Small-Molecule Activators of Protein Phosphatase 2A for the Treatment of Castration-Resistant Prostate Cancer. Cancer Res 78, 2065–2080, doi:10.1158/0008-5472.CAN-17-0123 (2018).

