## Supplementary Information for "Loss of PTPRB function remodels VEGFR1 activation in tumors overexpressing the receptor tyrosine kinase"

**Supplemental Data**

* Corresponding authors

Rahul Das:

**Keywords:** VEGFR, Receptor Tyrosine Kinase, cell signaling, Phosphatase, PTPRB.

### Methods

#### DNA constructs

The DNA constructs of VEGFR1 (Residue number 1–1338; pDONR223-FLT1 was a gift from William Hahn & David Root; Addgene plasmid # 23912)and VEGFR2 (Residue number 1-1356; pBE vector was a gift from Kalina Hristova Johns Hopkins University, Baltimore, MD; Addgene plasmid # 108854) fused to mCherry were cloned into pcDNA$^{TM}$3.1(+)[1]. The kinase dead VEGFR1 D1022N mutant was generated using PCR based methods. The VEGFR1 dimer and monomer controls were developed by replacing the transmembrane segment of VEGFR1 with the glycophorin A (GPA) transmembrane segment[1]. All the primers used to generate the DNA constructs are summarized in Table S1.

#### Cell culture

Chinese Hamster Ovary (CHO) cell line (obtained from National Centre for Cell Science- India) Human Embryonic Kidney 293T (HEK293T) cell line (obtained from ATCC) and M.D Anderson Metastatic Breast-231 (MDAMB231) cell line (obtained from National Centre for Cell Science- India) were cultured in Dulbecco's Modified Eagle Medium (DMEM) supplemented with 10% FBS, 100 units/mL penicillin, 100 µg/mL of streptomycin, and 0.25 µg/mL of amphotericin B at 37 °C with 95% humidity and 5% $CO_2$. Human Microvascular Endothelial Cell line-1 (HMEC-1) (obtained from ATCC) was cultured in Microcarrier Cell Development Buffer (MCDB-131) supplemented with 15% FBS, 100 units/mL penicillin, 100 µg/ml of streptomycin, 0.25 µg/ml of amphotericin B, 10mM glutamine, 1ng/ml hydrocortisone, and 10ng/ml EGF. Cells were tested for mycoplasma contamination before seeding. The cells at 50-60% confluency were transfected using Lipofectamine-3000 following the manufacturer's protocol. Plasmid DNA at a concentration of 500 ng/µl or 1000 ng/µl was used for microscopy or western blotting, respectively. The transfected cells were incubated for 14-16 hours in DMEM media supplemented with 10% FBS for protein expression. The cells were then serum-starved for six hours before being stimulated with 100 ng/ml $VEGF_{165}$. For imaging studies, the plasma membrane was labeled by treating the cells with WGA-Alexa Fluor 633 at a 1:200 dilution for 2 minutes. The excess dye was immediately removed by washing with 1x PBS.

#### Immunoblotting

The VEGFR1 constructs transiently expressed in CHO cells were activated with 100 ng/ml $VEGF_{165}$ or $Na_3VO_4$ at various concentrations. Kinase activity was quenched at the indicated time points (as indicated in each figure) by incubating the cells on ice. The cells were then washed with ice-cold 1X PBS and lysed by sonication after suspending in RIPA lysis buffer containing 10 mM Tris-Cl, pH 8.0, 140 mM NaCl, 1 mM EDTA, 0.1% SDS, and 1% Triton X-100 supplemented with protease inhibitors (2 mM Benzamidine, 1 mM PMSF) and 1mM $Na_3VO_4$. 50 µg of total protein was resolved on a 6% polyacrylamide gel and transferred to a PVDF membrane. VEGFR1 and phosphotyrosine 1213 (1213pY) were detected with the respective primary antibodies, listed in Table S2. The labeled primary antibody was visualized using an HRP-tagged secondary antibody. The blots were then developed with Western ECL substrate and imaged with a Chemidoc (Chemi XRQ GBOX DR4V2/2980). The expression of VEGFR1 and the extent of tyrosine 1213 phosphorylation were quantified by densitometric analysis of the blot performed using Fiji. Ver 1.54p[2]. Antibodies and materials used are listed in Table S2.

#### Quantitative RT-PCR

The expression level of VE-PTP (PTPRB) in HEK293T, HMEC-1, and MDAMB231 cell lines was measured using quantitative RT-PCR (qPCR). $1x10^6$ cells were washed with 1× PBS and lysed in TRIzol reagent ($1 \times 10^6$ cells). The lysate was mixed with chloroform (200 µL/mL), and centrifuged at 10,000 × g for 15 minutes at 4 °C. The aqueous phase containing the total RNA was collected, and the total RNA was precipitated by treating it with isopropanol overnight. The RNA in the pellet was collected by centrifugation at 10,000 × g for 10 minutes at 4 °C. The pellet was washed with 75% ethanol by centrifuging at 7,500 × g for 5 minutes, air-dried, and resuspended in RNase-free water. The cDNA was synthesized following the instructions of the Verso cDNA Synthesis Kit. The amount of VE-PTP cDNA was quantified by qPCR using gene-specific primers listed in Table S1 and SYBR Green Supermix. GAPDH was used as an internal control, and relative expression levels were calculated from the average Ct values of three biological and three technical replicates. Gene expression fold changes were calculated as: Fold change= $2^{-\Delta ct}$ where ΔCt = Ct (VE-PTP) – Ct (GAPDH)[3]. Relative mRNA level was plotted using GraphPad Prism 8.0.2. (Graphpad Inc., USA).

**Microscopy and image analysis**

**Immunofluorescence Confocal Microscopy**

Sample preparation: The CHO and HEK293T cell lines transiently expressing VEGFR2 and various constructs of VEGFR1 were treated with 100 ng/ml $VEGF_{165}$, $Na_3VO_4$, or Razuprotafib. The phosphorylation of Y residues at the C-terminal tail was quenched by immediately placing the cells on ice. The cells were washed with ice-cold 1X PBS and fixed with 4% PFA for 15 minutes. The cells were then washed twice with PBS before treating with 0.1M glycine for 15 minutes. The cells were permeabilized with 0.2% PBST (1X PBS and 0.2% Triton-X 100) for 5 minutes and incubated overnight at 4 °C with 3% BSA solution in 0.1% PBST (1X PBS and 0.1% Triton-X 100). The phosphorylation levels of 1213pY in VEGFR1 or 1175pY in VEGFR2 were determined using specific anti-phosphotyrosine antibodies, as listed in Table S2. Excess or unbound antibodies were removed by extensively washing with 0.1% PBST, followed by treatment with secondary antibody conjugated to FITC. Finally, cells were washed with 1X PBS before mounting on slides containing sufficient Prolong Gold antifade agent.

Imaging: A Leica SP8 confocal microscope was used to image fixed CHO and HEK293T cells. Images were acquired with a 63X oil-immersion objective with a pinhole set to 1 AU. All images were bidirectionally scanned in lightning mode at 700 Hz. Image resolution was set to 2048 x 2048 with the pixel size of 70nm. The mCherry-tagged VEGFR1 or VEGFR2 was imaged using a 40 mW, 552nm solid-state laser at 5% power with $\lambda_{ex}$ and $\lambda_{em}$ set at 552nm and 570-660nm, respectively. The FITC-tagged secondary antibodies were imaged using a 25 mW/488nm solid-state laser at 2% power with $\lambda_{ex}$ = 488nm and $\lambda_{em}$ = 500-550nm. WGA-tagged Alexa Fluor 633 was imaged with a 70 mW solid-state laser at 5% power with $\lambda_{ex}$ = 638 nm and $\lambda_{em}$ = 650-725nm, respectively.

Image analysis: The ligand-independent and ligand-dependent autophosphorylation of VEGF receptors was analyzed from the confocal images using Fiji. Ver 1.54p[2]. The ROI marking the plasma membrane or early endosome was identified from the Alexa Fluor 633 fluorescence[1,4]. The VEGFR expression and phosphorylation levels at the marked plasma membrane or in the early endosome were measured from the mean intensity ($\lambda_{em}$) of the red ($\lambda_{ex}$ = 552 nm) and green ($\lambda_{ex}$ = 488nm) channels, respectively. To study density-dependent receptor activation, the FITC intensity ($\lambda_{ex}$ = 488nm and $\lambda_{em}$ = 500-550nm) was plotted against the mCherry intensity ($\lambda_{ex}$ = 552nm and $\lambda_{em}$ =570-660nm). At each data point, cells were binned by mCherry intensity. The autophosphorylation kinetics of the VEGFR construct were measured using cells expressing intermediate levels of receptors (mCherry intensity of 1-2 x $10^4$ A.U). The ligand-dependent kinetics of VEGF receptors were determined from relative tyrosine phosphorylation levels:

$$Relative\ Phosphorylation = \frac{\frac{I_{t_n}^{FITC}}{I_{t_n}^{mCherry}} - \frac{I_{t_0}^{FITC}}{I_{t_0}^{mCherry}}}{\left(\frac{I_{t_n}^{FITC}}{I_{t_n}^{mCherry}}\right)_{max} - \frac{I_{t_0}^{FITC}}{I_{t_0}^{mCherry}}}$$

Where, $I_{t_n}^{FITC}$ is the FITC intensity at time t=n, $I_{t_n}^{mCherry}$ is the mCherry intensity at time t=n.

**Live Cell Imaging**

Sample preparation: The cells transiently expressing VEGFR constructs were serum starved for six hours and then transferred to HBSS buffer (136.893 mM NaCl, potassium chloride 5.366mM KCl, 261mM $CaCl_2$, 0.4057mM $MgSO_4.7H_2O$, 0.4919mM $MgCl_2.6H_2O$, 0.3371mM $Na_2HPO_4.2H_2O$ , 0.4409mM $KH_2PO_4$ , 5.551mM D-glucose and 4.166mM $NaHCO_3$). The cells were activated with 100 ng/ml $VEGF_{165}$, 0.1 mM $Na_3VO_4$, or 0.002 mM Razuprotafib, and the images was recorded.

Imaging: Live-cell imaging of CHO or HEK293T cells transiently expressing the VEGFR constructs was recorded using Abberior's Facility Line Stimulated Emission Depletion (STED) super-resolution microscope on an inverted Olympus IX83 body, with an Okolab stage-top incubator for $CO_2$ and temperature control. For 2D STED, the full-width half maximum (FWHM) was approximately 20 nm in the xy plane and ~500 nm along the z-axis achieved with a 775 nm pulsed STED depletion laser. The time-course videos of VEGFR-mCherry ($\lambda_{ex}$ = 561 nm) and WGA-Alexa Fluor 633 ($\lambda_{ex}$ = 640 nm) were acquired using a 60× objective (NA 1.42) and detected with an ultra-sensitive single-photon-counting APD. The dimerization and diffusion coefficients were quantified using 2D STED imaging of cells at the basal plane with the pixel size set at 20 nm and a dwell time of 5 µs. Image sequences were acquired over 300 frames at 4 ms intervals using 15% and 5% laser power for the 561 nm and 640 nm channels, respectively, with accumulation set to 1. Step-wise photobleaching was carried out with a pixel size of 50 nm, using 50 pre-bleach frames at 15% (561 nm) and 5% (640 nm) laser power, followed by 400 bleach frames at 100% laser power and 5 post-bleach frames acquired under pre-bleach laser settings. All images were acquired using a 1 AU pinhole.

Image analysis: All image and data analysis tasks were performed in MATLAB R2025a. All code and software packages compatible with MATLAB R2025a on a Linux 64-bit operating system were obtained from GitHub[5]. STED microscopy videos (obf files) were loaded into MATLAB using Bio-Formats reader. Particle detection and tracking were performed using u-track[5]. The detection (of particle) and tracking parameters were optimized, as listed in Table S3. Intensity distributions were represented as probability density functions obtained by normalizing the corresponding frequency histograms to unit area[6,7] as shown in the following equations.

$$1 = \sum_{i=imin}^{imax} p(i)\Delta \mathrm{i}$$

$$p(i) = \frac{\text{Counts in bin}}{\mathrm{N}\Delta\mathrm{i}}$$

Where, N = Total number of particles, Δi = bin width (from histogram) , *p(i)* = probability density at intensity bin i, $i_{min}$ and $i_{max}$ define the full intensity range of the histogram.

The normalization does not alter the shape of the intensity distribution, facilitating quantitative comparisons and subsequent fitting with Gaussian models[8]. A mixed Gaussian fit was performed on the distribution of particle intensities to estimate the fraction of total particles in each underlying component. The mixed Gaussian fitting on the intensity histogram is defined with the equation below –

$$\varphi(i) = \sum_{n=1}^{n_{max}} w_n \cdot \frac{1}{\sigma\sqrt{2\pi}} e^{-\frac{(i-\mu\cdot n)^2}{2(\sigma\cdot n)^2}}$$

Where *φ(i)* is the probability density of particles having intensity i, n is the component number, and $w_n$ is the fraction of area under the curve of component n ($\sum w_n = 1$). μ and σ are the mean and Standard deviation of reference single fluorophores respectively. The maximal number of components ($n_{max}$) was determined for each case by progressively increasing $n_{max}$ until the addition of one component no longer resulted in a statistically better fitting, as judged by an F-test ($P > 0.05$). For our case, the $n_{max}$ value was set at 2. Based on the u-track data, particle density ranges were optimized to determine a linear particle density range (0.15–0.25 particles/$\mu m^2$) (Figure S3). Beyond this density range, the monomer and dimer control showed spontaneous oligomerization. Therefore, in all our live-cell experiments to detect receptor dimerization, a particle density range of 0.15–0.25 particles/$\mu m^2$ was used. Additionally, the oligomeric size of the particle was determined from stepwise photobleaching of the particle intensity[9]. A plot of mean-squared displacement (MSD) versus time was generated from the u-Track particle tracking data. The effective diffusion coefficient (*D*) of each valid track was calculated from the mean square displacement ($r^2$),

$$D = \frac{r^2}{4\Delta t}$$

where $\Delta t$ is the time for which the MSD vs time plot shows a linear fit. The mean diffusion coefficient is proportional to the slope of the MSD vs time plot (Figure S4). The probability density function for the diffusion coefficient was calculated as above.

**Transcriptome analysis**

The mRNAseq data for the cancer patients deposited in the TCGA databank[10] (which was accessed through UCSC Xena) were used for this study[11]. Pan-cancer mRNA expression of VEGFR1, VEGFR2, and VEGF-A was compared with the mRNA expression in the adjacent normal tissue[10]. For statistical significance, samples with $n \geq 50$ were analyzed using the student's t-test. (see Table S4). The survival analysis of KIRC patients was performed using the survival data obtained from UCSC Xena. The tumor samples were divided into two categories: tumor alive and tumor dead. The Kaplan-Meier curve and survival percentage were plotted for high and low PTPRB expression. All plots were generated using Python 3.13, and the following packages were used: pandas, numpy, matplotlib, and lifelines (KaplanMeierFitter)[12].

**Proteome and phosphoproteome analysis of KIRC**

Proteomic and phosphoproteomic data for VEGFR1 and VEGFR2 in KIRC patients were obtained from the Clinical Proteomic Tumor Analysis Consortium (CPTAC)[13] and accessed through LinkedOmics[14]. The protein expression or phosphorylation levels of VEGFR1 and VEGFR2 in cancer tissue were compared with those in the adjacent normal tissue of a KIRC patient[13]. For statistical significance, samples with $n \geq 50$ were considered and analyzed using the student's t-test.

**Mutation analysis**

Mutation frequencies for EGFR, VEGFR1, and VEGFR2 in pan-cancer and KIRC patients were determined from data deposited in COSMIC[15] and the cBioPortal repository[16], respectively. The occurrence of a residue-specific missense mutation in each receptor was presented as a percentage of the total sample. For pan-cancer data, mutation frequencies of <0.02%, 0.02-0.6%, and >0.6% for a particular residue are defined as low, medium, and high, respectively.

**Gene network analysis**

The GENIE3 method was applied on RNAseq - IlluminaHiSeq dataset of KIRC patients (TCGA KIRC database, Id - TCGA.KIRC.sampleMap/HiSeqV2). The dataset was downloaded from UCSC XENA[11]. Using TCGA sample codes, the samples were separated into two groups: normal (72 samples) and tumor (533 samples)[10]. The GENIE3 code is available on GitHub[17]. The code was run using Python 3.13 on a 64-bit Linux system with 256 GB RAM and an Intel Xeon Silver 2.1 GHz processor. The parameters used are listed in Table S4. A pair of gene regulatory networks (GRNs) was inferred using GENIE3, one from normal samples and the other from tumor samples. GENIE3 assigned a confidence score between 0 and 1 to each edge. It represents GENIE3's confidence in the presence of that edge. For each GRN, the subnetwork in which VEGFR1 was either the regulator or the target gene was extracted. The subnetwork was refined by retaining only the top 50 edges based on their confidence scores. A second set of subnetworks was generated by extracting the top 50 edges where VEGFR1 is either the regulator or the target of a receptor tyrosine phosphatase. For network analysis, several R packages were used (dplyr, igraph, tidygraph, ggraph, ggplot2 among the major ones). The R source code was written using RStudio 2025.09.2 Build 418.

**Statistics and reproducibility**

The results have been expressed as the mean ± SD from more than three replicates ($n \geq 3$). All statistical analyses were done using Graphpad Prism 8.0.2 software (Graphpad Inc., USA). Comparison between two groups was made using Student's t-test. Statistical significance was assessed using the following criteria: $^{*}p < 0.05$, $^{**}p < 0.01$, $^{***}p < 0.001$, $^{****}p < 0.0001$, ns denotes not significant.

### Supplementary Tables

**Supplementary Table S1: List of Primers**

| | Primer | Sequence |
|---|---|---|
| VEGFR1 - D1022N | Fwd | GTG CAT TCA TCG GAA CCT GGC AGC GAG |
| | Rev | CTCGCTGCCAGGTTCCGATGAATGCAC |
| VEGFR1 - GPA | GPA_Fwd 1 | TACCGGACTCAGATCTCGAGATGGTCAGCTACTGGGACACCGG |
| | GPA_Rev 1 | CTCCAGATTAGACTTGTCCGAG |
| | GPA_Fwd 2 | CGGACAAGTCTAATCTGGAGCTCATTATTTTGGGGTGATGG |
| | GPA_Rev 2 | ACCGTAAGAAATTAAGAGGATCGTTCCAATAACACC |
| | GPA_Fwd 3 | TCCTCTTAATTTCTTACGGTATC CGA AAA ATG AAA AGG TCT TC |
| | GPA_Rev 3 | GGTGGCGACCGGTGGATCCACGATGGGTGGGGTGGAGTACAGGA |
| VEGFR1 - G83I | Fwd | GTGATGGCTATTGTTATTGGAACG |
| | Rev | CCCAAAAATAATGAGCTCCAG |
| PTPRB-Human | Fwd | GGGCTCACCCTGTAACTTTAGC |
| | Rev | TCTATCCGAAAGGTAGGGCAC |
| GAPDH-Human | Fwd | TGTGGGCATCAATGGATTTGG |
| | Rev | ACACCATGTATTCCGGGTCAAT |

**Supplementary Table S2: List of antibodies and dyes**

| Antibodies | Source | Details | Dilution |
|---|---|---|---|
| Phospho-VEGF Receptor 1 (Tyr1213) Polyclonal Antibody Rabbit | Thermofisher Scientific | Cat # PA5-99362 79445753 | 1:1000 (IB) 1:200 (IF) |
| Phospho-VEGF Receptor 2 (Tyr1175) Rabbit monoclonal antibody (19A10) | Cell Signaling Technology, (Danvers, MA, USA) | Cat # 2478T Lot:17 | 1:200(for IF) |
| VEGFR1 Goat polyclonal antibody | R & D system (Minneapolis, MN, USA) | Cat # AF321 Lot: AHT2018011 | 1:1000 (for IB) |
| Rabbit HRP Secondary antibody | Abcam (Waltham, MA 02453, USA) | Cat# 50095 Lot: 2960660 | 1:2000 (for IB) |
| Secondary Alexa Fluor™ 488 | Thermofisher Scientific | Cat # A-11008 | 1:500 (IF) |
| Wheat Germ Agglutinin (WGA)-633 | Thermofisher Scientific | Cat #W21404 Lot: 3328748 | 1:200 (IF & Live cell) |

**Supplementary Table S3: Parameters for u-track analysis**

| Parameters | VEGFR-mCherry channel |
|---|---|
| Gaussian standard deviation | 1 pixel |
| Pixel size | 50 nm |
| Local maxima detection: Use rolling window time-averaging | No |
| Local maxima detection: α-value for comparison with local background | 0.5 |
| Gaussian mixture-model fitting at local maxima: Do iterative Gaussian mixture model fitting | Yes |
| Gaussian mixture-model fitting at local maxima: α-value for residuals test | 0.05 |
| Gaussian mixture-model fitting at local maxima: α-value for amplitude test | 0.05 |

| | |
|---|---|
| Gaussian mixture-model fitting at local maxima: α-value for distance test | 0.9 |
| Overall: Do segment merging | Yes |
| Overall: Do segment splitting | Yes |
| Frame-to-frame linking: Brownian search radius upper bound | 5 pixels |
| Frame-to-frame linking: Multiplication factor for Brownian search radius calculation | 3 |
| Gap closing, merging and splitting – general: Brownian search radius upper bound | 5 pixels |
| Gap closing, merging and splitting – general: Multiplication factor for Brownian search radius calculation | 3 |
| Gap closing, merging and splitting – general: Scaling power to expand Brownian search radius | 0.5 |
| Gap closing, merging and splitting – general: Use nearest neighbor distance to expand Brownian search radius | Yes (5 frames for nearest neighbor calculation) |
| Gap closing, merging and splitting – merging & splitting: Search radius lower bound | 2 pixels |
| Gap closing, merging and splitting – merging & splitting: For an end/start, the possibility of merging/splitting is allowed only if there is no possibility of gap closing | No |
| Gap closing, merging and splitting – birth & death: Birth and death cost | Percentile (Automatic) |

**Supplementary Table S4: Pan cancer mRNA expression deposited in the UCSC Xena.**

| Cancer | VEGFR2<br>Normal log2RSEM [n]<br>Tumor log2RSEM [n] | p value | VEGFA<br>Normal log2RSEM [n]<br>Tumor log2RSEM [n] | p value | VEGFR1<br>Normal log2RSEM [n]<br>Tumor log2RSEM [n] | p value |
|---|---|---|---|---|---|---|
| BLCA | 9.1±1.0[19]<br>8.3±1.1[407] | 1.5E-03 | 11.5±1.3[19]<br>11.9±1.2[407] | 1.1E-01 | 9.6±0.9[19]<br>9.3±1.0[407] | 9.2E-02 |
| BRCA | 10.8±0.7[114]<br>9.6±0.9[1097] | <1E-4 | 10.3±0.5[114]<br>10.8±0.9[1097] | <1E-4 | 10.7±0.7[114]<br>9.9±0.8[1097] | <1E-4 |
| CESC | 12.1±1.8[3]<br>7.3±1.2[303] | <1E-4 | 12.2±1.0[3]<br>11.7±1.0[303] | 3.3E-01 | 11.1±0.8[3]<br>8.6±1.0[303] | <1E-4 |
| CHOL | 10.3±0.8[9]<br>8.7±1.1[36] | 9.5E-04 | 12.0±0.4[9]<br>12.3±1.0[36] | 2.5E-01 | 9.4±0.7[9]<br>9.6±0.8[36] | 1.8E-01 |
| COAD | 8.4±0.6[41]<br>8.4±1.0[286] | 7.8E-01 | 10.3±0.6[41]<br>11.6±0.7[286] | <1E-4 | 8.8±0.6[41]<br>9.2±0.9[286] | 1.2E-02 |
| COAD READ | 8.4±0.6[51]<br>8.5±1.0[380] | 6.4E-01 | 10.3±0.6[51]<br>11.6±0.7[380] | <1E-4 | 8.8±0.7[51]<br>9.2±0.9[380] | 6.0E-04 |
| ESCA | 9.5±1.1[11]<br>8.7±1.2[184] | 5.6E-02 | 11.4±0.5[11]<br>11.5±1.0[184] | 1.8E-01 | 10.5±1.4[11]<br>9.7±1.1[184] | 3.5E-01 |
| GBM | 7.6±0.6[5]<br>9.0±0.8[154] | 7.0E-04 | 9.7±0.5[5]<br>12.6±1.6[154] | 3.4E-04 | 10.1±0.6[5]<br>9.6±0.8[154] | 3.3E-01 |
| GBMLGG | 7.6±0.6[5]<br>8.7±0.9[670] | 2.5E-02 | 9.7±0.5[5]<br>9.5±1.8[670] | 7.2E-01 | 10.1±0.6[5]<br>9.8±0.7[670] | 7.0E-01 |
| HNSC | 8.0±1.3[44]<br>8.3±1.2[520] | 9.3E-01 | 10.5±0.7[44]<br>11.3±1.0[520] | <1E-4 | 8.4±1.1[44]<br>9.3±0.9[520] | <1E-4 |
| KICH | 10.6±0.6[25]<br>10.2±1.0[66] | 1.2E-01 | 11.5±0.7[25]<br>12.3±0.8[66] | <1E-4 | 10.8±0.5[25]<br>10.9±1.1[66] | 8.4E-01 |
| KIRC | 11.1±0.8[72]<br>12.3±1.4[533] | <1E-4 | 11.5±0.7[72]<br>14.9±1.2[533] | <1E-4 | 11.5±0.8[72]<br>13.5±1.3[533] | <1E-4 |
| KIRP | 10.9±0.7[32]<br>7.1±1.5[290] | <1E-4 | 11.6±0.6[32]<br>9.8±1.4[290] | <1E-4 | 11.3±0.7[32]<br>8.5±1.2[290] | <1E-4 |

| | | | | | | |
|---|---|---|---|---|---|---|
| LIHC | 10.1±0.8[50]<br>9.3±1.3[371] | <1E-4 | 11.7±0.5[50]<br>11.7±0.8[371] | 6.5E-01 | 9.3±0.8[50]<br>9.7±0.9[371] | 5.4E-03 |
| LUAD | 11.3±0.7[59]<br>10.0±1.3[515] | <1E-4 | 12.2±0.5[59]<br>12.2±1.0[515] | 2.9E-01 | 10.7±0.7[59]<br>10.1±0.9[515] | <1E-4 |
| LUNG | 11.3±0.7[110]<br>9.1±1.4[1017] | <1E-4 | 12.1±0.5[110]<br>12.0±1.0[1017] | 5.3E-01 | 10.8±0.7[110]<br>9.8±0.9[1017] | <1E-4 |
| LUSC | 11.3±0.7[51]<br>8.5±1.1[502] | <1E-4 | 12.0±0.5[51]<br>11.9±0.9[502] | 8.7E-01 | 10.8±0.6[51]<br>9.6±0.8[502] | <1E-4 |
| PAAD | 9.4±0.3[4]<br>9.2±1.0[178] | 5.5E-01 | 11.4±0.7[4]<br>11.8±0.9[178] | 7.7E-01 | 10.2±0.5[4]<br>10.3±0.8[178] | 7.5E-01 |
| PCPG | 10.2±0.2[3]<br>10.4±1.3[179] | 9.2E-01 | 11.8±0.6[3]<br>11.8±1.2[179] | 9.0E-01 | 10.3±0.2[3]<br>11.1±1.3[179] | 2.2E-01 |
| PRAD | 9.8±1.0[52]<br>8.9±1.0[497] | <1E-4 | 12.4±1.4[52]<br>11.6±1.5[497] | 2.8E-03 | 9.7±0.9[52]<br>9.4±1.1[497] | 5.0E-02 |
| READ | 8.3±0.7[10]<br>8.8±0.9[94] | 1.7E-01 | 9.9±0.7[10]<br>11.6±0.7[94] | <1E-4 | 8.9±0.8[10]<br>9.4±0.8[94] | 1.1E-02 |
| SARC | 10.3±2.0[2]<br>9.8±1.2[259] | 5.2E-01 | 10.8±1.9[2]<br>10.9±1.3[259] | 7.8E-01 | 10.4±1.9[2]<br>10.2±1.1[259] | 8.8E-01 |
| SKCM | 7.7±0.0[1]<br>8.2±1.1[104] | NA | 9.0±0.0[1]<br>9.6±1.4[104] | NA | 8.5±0.0[1]<br>8.2±1.1[104] | NA |
| THCA | 10.7±0.6[59]<br>10.7±1.1[505] | 9.1E-01 | 13.2±0.6[59]<br>12.7±1.0[505] | <1E-4 | 11.1±0.6[59]<br>11.4±1.1[505] | 1.1E-01 |
| THYM | 8.9±1.5[2]<br>8.2±1.4[120] | 4.1E-01 | 8.9±1.4[2]<br>9.4±1.5[120] | 6.9E-01 | 9.1±2.0[2]<br>8.0±1.2[120] | 2.7E-01 |
| UCEC | 11.2±1.5[24]<br>8.0±1.1[176] | <1E-4 | 11.4±1.2[24]<br>11.4±1.0[176] | 1.1E-01 | 10.2±1.6[24]<br>9.3±0.9[176] | <1E-4 |

**Supplementary Table S5: Parameters for VEGFR1 network analysis**

| **Parameter** | **Setting** |
|---|---|
| Regulators | all |
| Regression Method | RF |
| K | sqrt |
| ntrees | 500 |

**Supplementary Table S6: Mean intensity of VEGFR-mCherry constructs from 2D STED live cell imaging.**

| Construct/Condition | Mean Intensity of monomer component (AU) | Mean Intensity of dimer component (AU) | Overall mean intensity (AU) |
|---|---|---|---|
| VEGFR1-G83I | 2.6719 ± 1.8899 | - | 2.6719 |
| VEGFR1-GPA | - | 6.1452 ± 2.5535 | 6.1452 |
| VEGFR1 | 3.0023 ± 1.4704 | 6.0362 ± 2.0553 | 3.1583 |
| VEGFR1 + 100 ng/mL $VEGF_{165}$ | 2.9921 ± 1.9619 | 6.1328 ± 2.0192 | 3.5295 |
| VEGFR1 + 0.1mM $Na_3VO_4$ | 3.0107 ± 1.5947 | 5.9929 ± 2.2132 | 3.7976 |
| VEGFR1 + 0.002 mM Razuprotafib | 3.0019 ± 1.9188 | 6.0054 ± 2.1922 | 3.7504 |
| VEGFR2 | 2.8743 ± 1.9004 | 6.1114 ± 2.1623 | 3.4603 |
| VEGFR2 + 100 ng/mL $VEGF_{165}$ | 3.1864 ± 1.9641 | 6.1478 ± 2.2232 | 3.9269 |
| VEGFR2 + 0.1mM $Na_3VO_4$ | 3.2232 ± 1.6516 | 6.1354 ± 2.2377 | 4.0637 |

### Supplementary figures

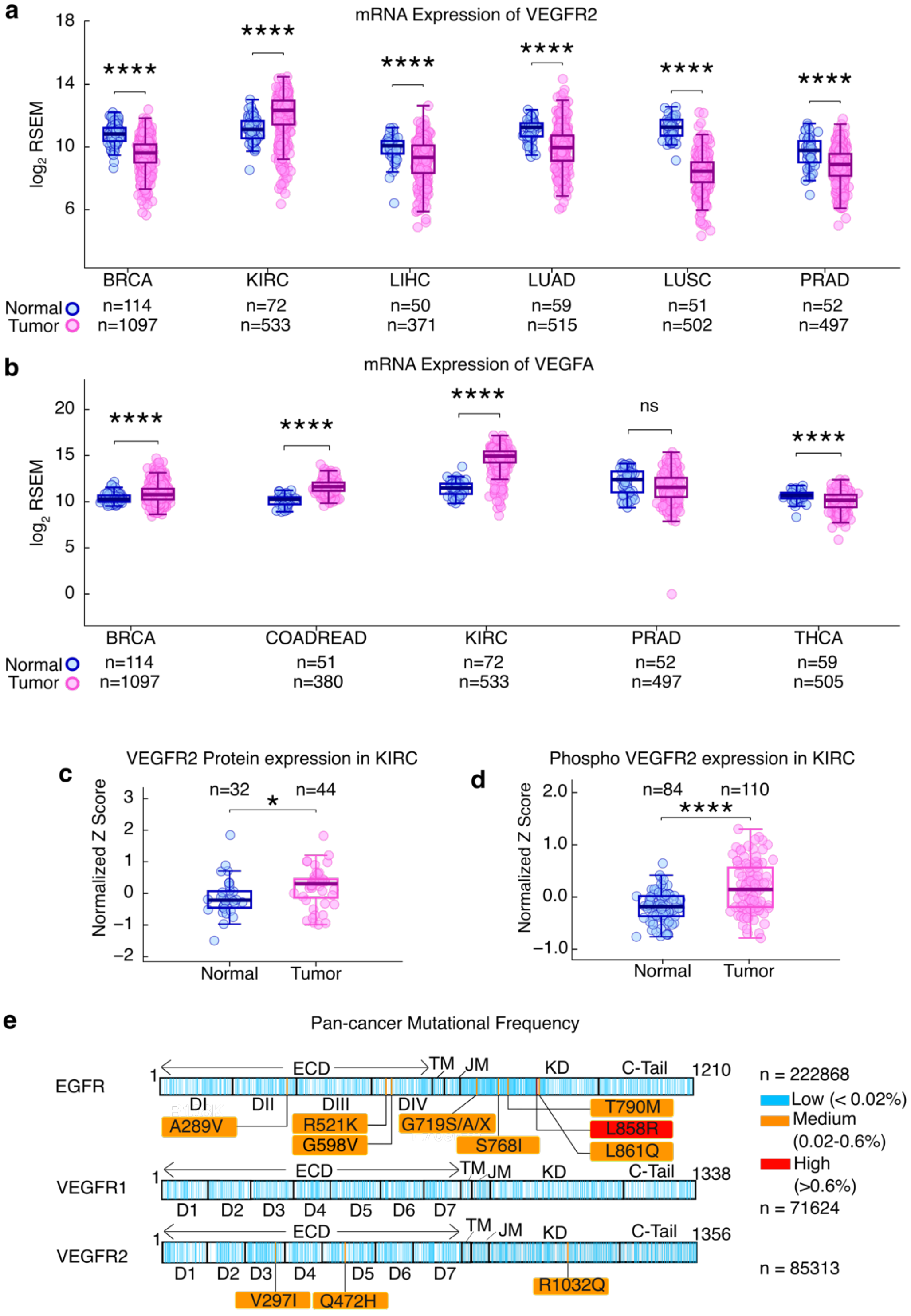

**Figure S1: Multi-omics analysis of VEGFR2 and VEGFA in cancer patients**

**a-b)** The normalized mRNA level ($log_2$RSEM) of VEGFR2 (panel a) and VEGFA (panel b) is plotted for normal tissue (blue) and tumor tissue (pink) from the indicated cancer patients. The patient information was retrieved from the TCGA-curated cancer dataset.

**c)** Plot of normalized VEGFR2 protein expression level (normalized Z score) in normal tissue (blue) and tumor tissue (pink) from KIRC patients.

**d)** Plot of normalized VEGFR2 phosphorylation level in normal (blue) and tumor (pink) tissue from KIRC patients.

**e)** Pan-cancer mutation frequency in EGFR, VEGFR1, and VEGFR2 reported in COSMIC data bank is mapped on the secondary structure of the receptor kinases. The region where no mutations were detected is shown in white. The vertical color lines indicate the mutation frequency of the amino acid residue. For EGFR, VEGFR1 and VEGFR2 the number of patient samples was 222868, 71624 and 85313, respectively.

BRCA- Breast Cancer, COADREAD- Colonic and rectal adenocarcinoma, KIRC- Kidney renal clear cell carcinoma, LIHC- Liver hepatocellular carcinoma, LUAD- Lung adenocarcinoma, LUSC-lung squamous cell carcinoma, PRAD- Prostate adenocarcinoma, THCA- Thyroid carcinoma

In panels **a-d)**, each circle represents a patient. The solid horizontal line is the median, and the error indicates ±SD. Comparison between the two groups was made using Student's t-test. Statistical significance was assessed using the following criteria: $^{*}p< 0.05$, $^{**}p< 0.01$, $^{***}p< 0.001$, $^{****}p< 0.0001$, and ns denotes not significant.

**e)** The schematic is made using Inkscape Ver1.4.3.

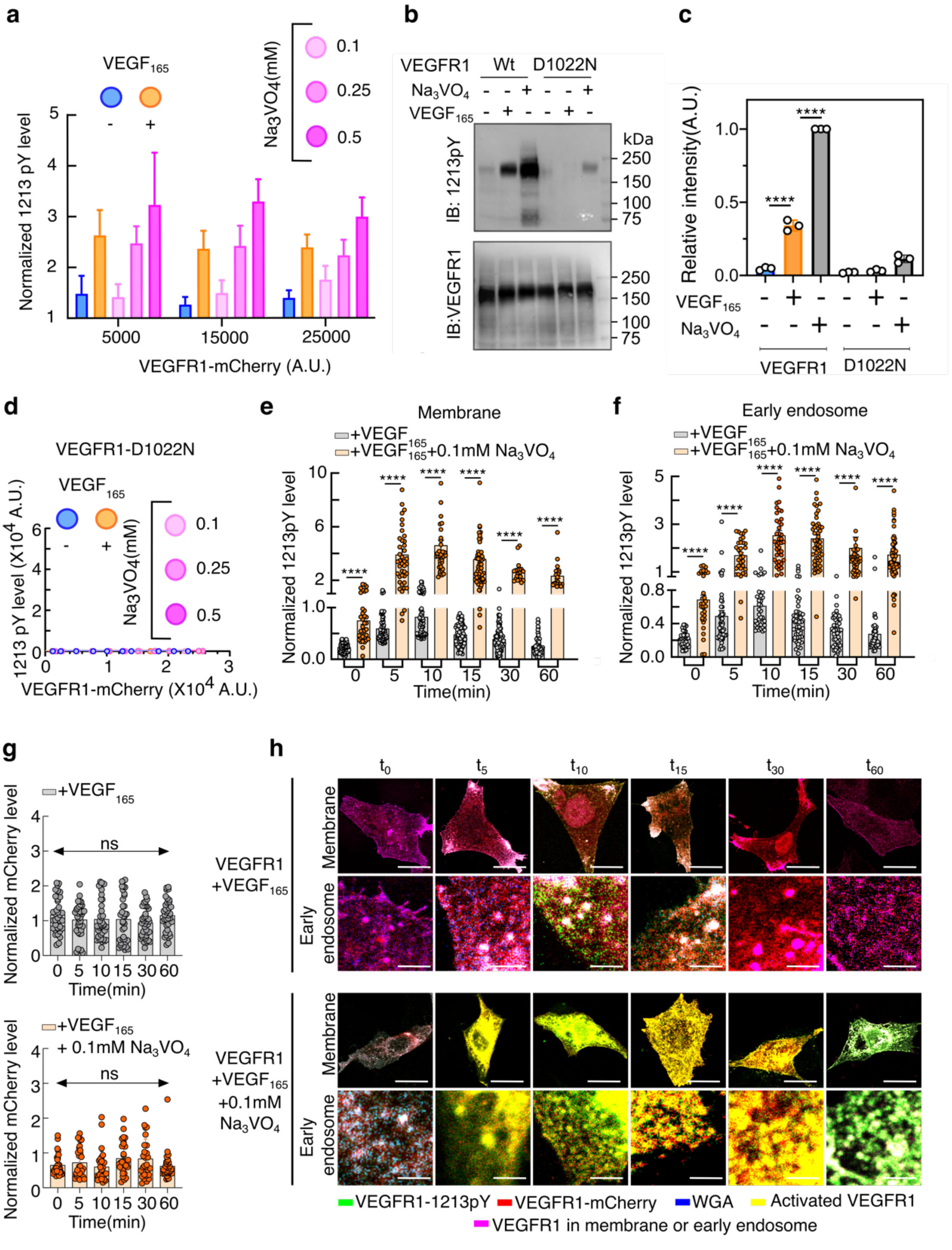


**Figure S2: Effect of Pan-phosphatase inhibitor, $Na_3VO_4$, on ligand-independent and dependent VEGFR1 autophosphorylation**

**a)** Concentration-dependent phosphorylation of 1213pY at the indicated sodium orthovanadate ($Na_3VO_4$) concentration is plotted against the VEGFR1 expression level.

**b)** Representative immunoblot of VEGFR1 and VEGFR1 Kinase dead mutant (D1022N) phosphorylation by 100ng/ml $VEGF_{165}$ and 0.5mM of $Na_3VO_4$.

**c)** Densitometric analysis of the immunoblot in panel b. The 1213pY intensity was normalized by the VEGFR1 loading control.

**d)** The VEGFR1(D1022N)-mCherry intensity at the plasma membrane is plotted against FITC intensity, denoting 1213pY level, n=100-120 cells. At each data point, cells were binned based on mCherry intensity. Each data point represents the mean ± SD from approximately 10–20 cells. The solid lines are guiding lines. The VEGFR1 was activated either with 100ng/ml of $VEGF_{165}$ or at the indicated concentration of $Na_3VO_4$.

**e-f)** Bar plot of normalized 1213pY levels at the plasma membrane (panel e) or the early endosome (panel f) against time. The FITC intensity corresponding to 1213pY levels was normalized against the VEGFR1-mCherry intensity. Comparison between the two groups was made using Student's t-test. Statistical significance was assessed using the following criteria: $^{*}p< 0.05$, $^{**}p< 0.01$, $^{***}p< 0.001$, $^{****}p< 0.0001$, ns denotes not significant.

**g)** Bar plot of normalized VEGFR1 intensity at the plasma membrane against time. The VEGFR1-mCherry constructs were transiently expressed in CHO cells and stimulated with 100ng/ml $VEGF_{165}$ (upper panel) or 100ng/ml $VEGF_{165}$ in the presence of 0.1mM of $Na_3VO_4$ (lower panel). The mCherry intensity corresponding to VEGFR1 levels was normalized against the intensity of the membrane marker WGA-Alexa Fluor 633. Comparison between the groups was performed using Student's t-test. Statistical significance was assessed using the following criteria: *p< 0.05, **p< 0.01, ***p< 0.001, ****p< 0.0001, ns denotes not significant.

**h)** Representative time-resolved confocal images of CHO cells, transiently expressing VEGFR1, following stimulation with 100ng/ml $VEGF_{165}$ (row 1 and 2) or 100ng/ml $VEGF_{165}$ in the presence of 0.1mM of $Na_3VO_4$ treatment (row 3 and 4). The images in rows 1 and 3 are of whole cells. Rows 2 and 4 are the images of the early endosome. The red channel ($\lambda_{ex}$ =552nm, $\lambda_{em}$ = 570-660nm), which denotes VEGFR1 expression. The green channel ($\lambda_{ex}$ = 488nm, $\lambda_{em}$ = 500-550nm) denotes Y1213 phosphorylation level (1213pY). The blue channel ($\lambda_{ex}$ = 638 nm and $\lambda_{em}$ = 650-725nm) denotes plasma membrane and early endosomes labeled with WGA-tagged Alexa Fluor 633. The yellow and magenta represent the merged images. Cells with mCherry intensity of 1-2 x $10^4$ AU at the plasma membrane were considered in this study. For rows 1 and 3, Scale bar=10 µm; For rows 2 and 4, Scale bar=1 µm.

**a, c-f)** The plots are made using Graphpad prism 8.0.2.

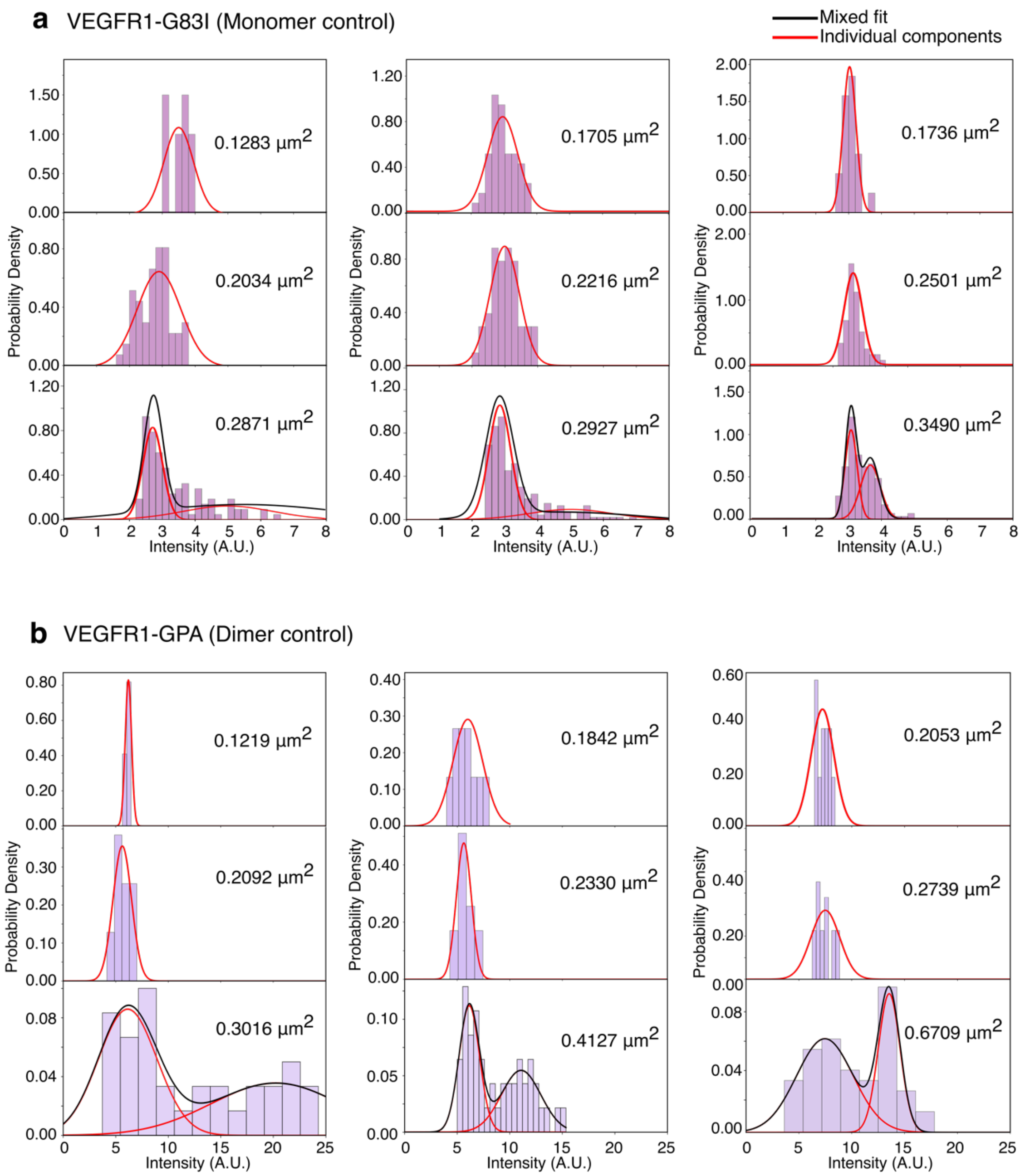


**Figure S3 : Optimization of linear density range for quantitative measurement of VEGFR1 oligomers**

The density of each protein cluster was estimated based on the number of particles within that cluster that form valid tracks.

**a-b)** Histogram plot of intensity distribution versus probability density for the monomer control (panel a) and dimer control (panel b). Each panel represents the intensity measured for all particles in n=12-20 cells at the indicated densities. The red and black lines show single and mixed Gaussian fits, respectively. The plots are made using MATLAB R2026a.

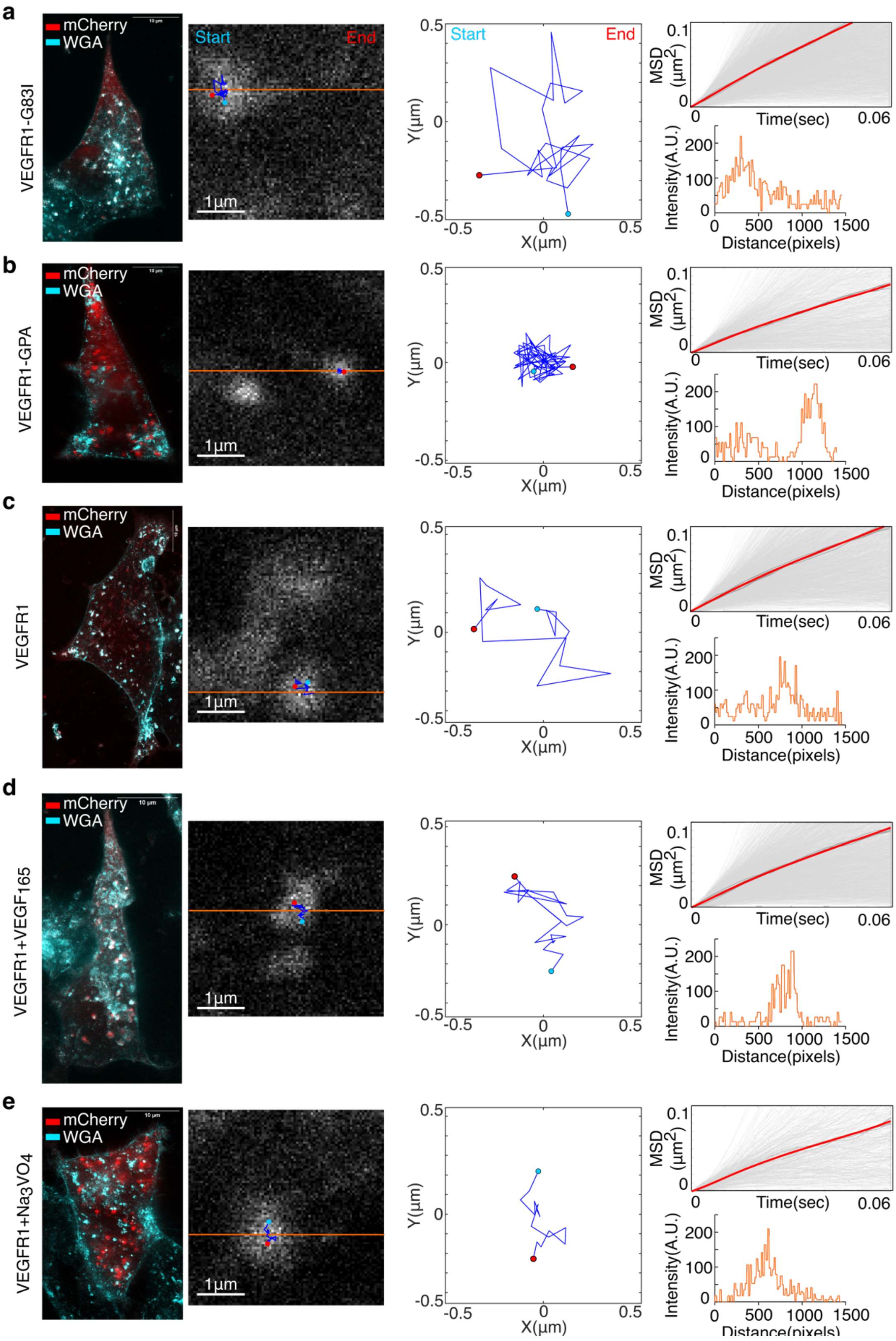


**Figure S4 : Particle mobility and trajectory analysis of individual VEGFR1 particles.**

**a-e)** From left to right in each panel, are the representative whole-cell 2D STED images at the basal plane of CHO cells transiently expressing the indicated VEGFR1 construct (n=12-20 cells). The

VEGFR1-GPA and G83I mutant of VEGFR1-GPA served as dimer and monomer controls, respectively. The cells were stimulated with 100 ng/ml $VEGF_{165}$ or 0.1 mM $Na_3VO_4$.

The second column from the left shows a 4 µm x 4 µm representative region in the cell, visualizing a cluster considered for particle mobility analysis. The particle trajectory is shown in a blue line, the start and end points marked in cyan and red, respectively.

The third column from the left shows the same particle track in a 1 µm x 1 µm region, with a white background for clarity. The plot is made using MATLAB R2026a.

The fourth column from the left is the Mean Squared Displacement (MSD) of all individual particle tracks, plotted against time (0 to 0.06s). The individual tracks are shown in grey, and the mean is shown in red. The plot is made using MATLAB R2026a.

The bottom panel on the right shows the pixel intensity on the orange line drawn in the representative whole-cell 2D STED image. This plot is made using Fiji ImageJ 1.54p.

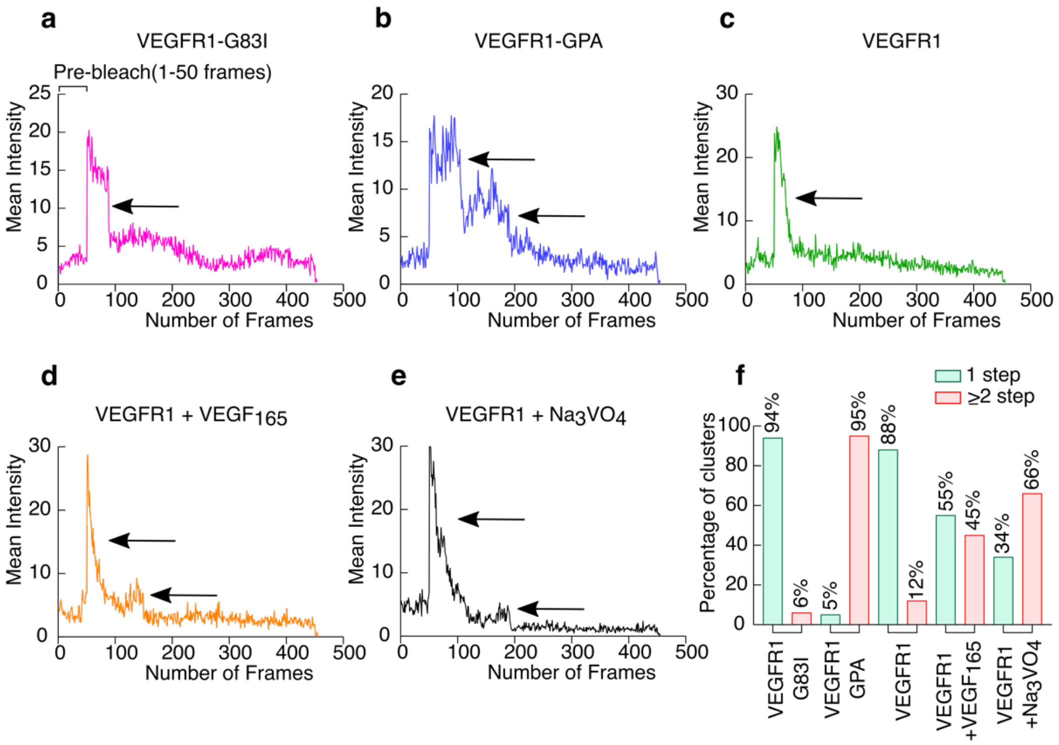


**Figure S5: Oligomeric distribution of VEGFR1**

**a-e**) Representative stepwise photobleaching traces of VEGFR1 clusters, plotted as mean intensity versus frame number. For mCherry bleaching, the laser power was set to 100%, and 400 frames were acquired every 4ms. 50 pre-bleach images and 5 post-bleach images were recorded. The arrow points to the distinct bleaching steps of the individual cluster.

**f)** Bar plot representing the percentage of clusters showing one step (green bar) or ≥ two steps (red bar) bleaching for the indicated VEGFR1 constructs. CHO cells transiently expressing VEGFR1-G83I (monomer control) , VEGFR1-GPA (dimer control), and VEGFR1 were activated with 100ng/ml $VEGF_{165}$ or 0.1mM $Na_3VO_4$ (n=15-25 cells per condition).

**a-f)** The plots are made using Graphpad prism 8.0.2.

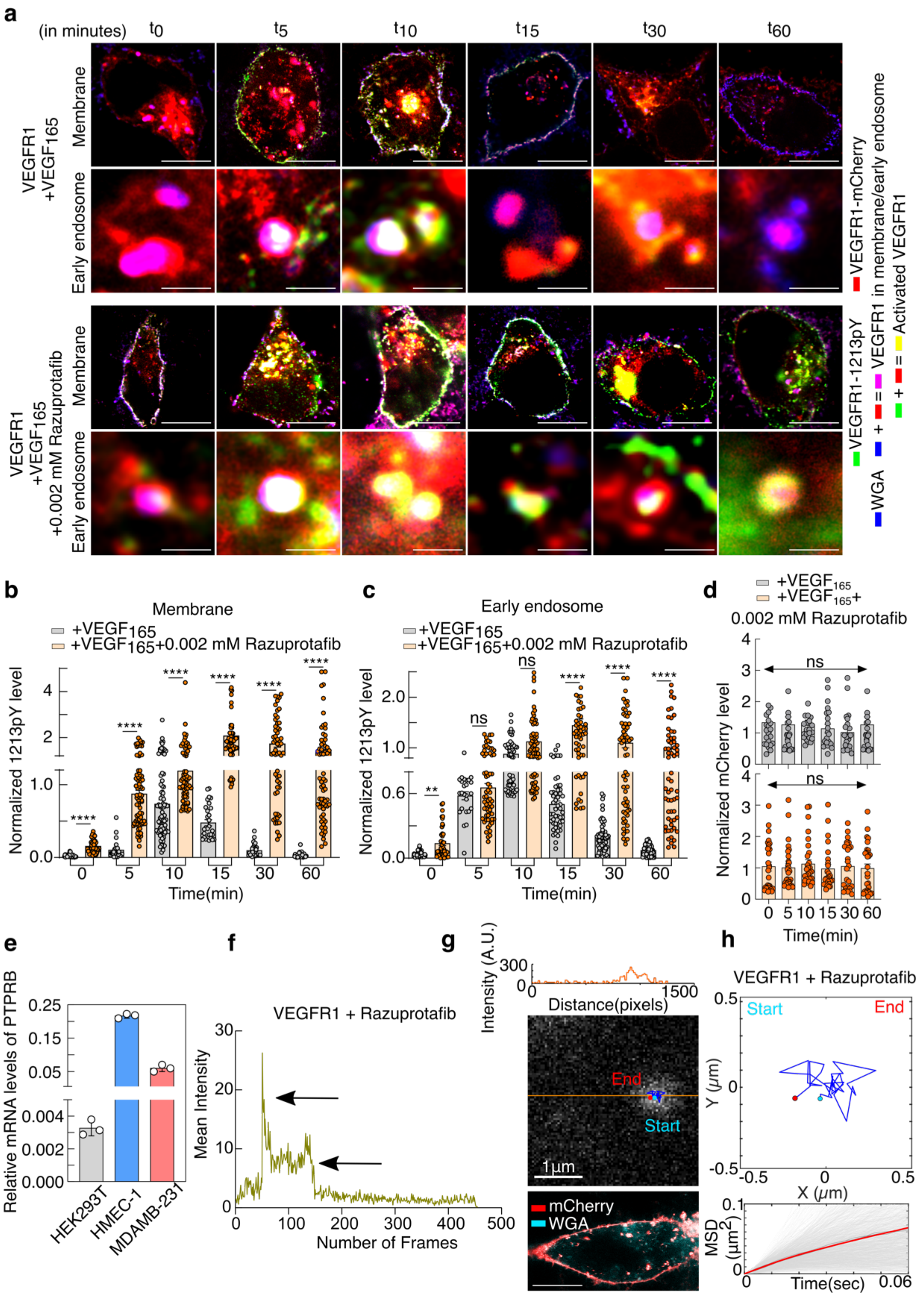


**Figure S6: Effect of Razuprotafib on VEGFR1 phosphorylation kinetics and oligomerization.**

**a)** Representative time-resolved confocal images of transiently VEGFR1 expressing HEK293T cells following stimulation with 100ng/ml $VEGF_{165}$ (row 1 and 2) and 100ng/ml $VEGF_{165}$ in presence of 0.002mM Razuprotafib treatment (row 3 and 4). The images in rows 1 and 3 are of whole cells. Rows

2 and 4 are the images of the early endosome. The red channel ($\lambda_{ex}$ =552nm, $\lambda_{em}$ = 570-660nm), which denotes VEGFR1 expression. The green channel ($\lambda_{ex}$ = 488nm, $\lambda_{em}$ = 500-550nm) denotes Y1213 phosphorylation level (1213pY). The blue channel ($\lambda_{ex}$ = 638 nm and $\lambda_{em}$ = 650-725nm) denotes the plasma membrane and early endosomes labeled with WGA-tagged Alexa Fluor 633. The yellow and magenta represent the merged images. Cells expressing with mCherry intensity of 1-2 x $10^4$ A.U at the plasma membrane were considered for this study. For rows 1 and 3, Scale bar=10 µm; For rows 2 and 4, Scale bar=500 nm

**b-c)** Bar plot of normalized 1213pY levels at the plasma membrane (panel b) or the early endosome (panel c) against time. The FITC intensity corresponding to 1213pY levels was normalized against the VEGFR1-mCherry intensity. Comparison between the two groups was made using Student's t-test. Statistical significance was assessed using the following criteria: $^{*}p< 0.05$, $^{**}p< 0.01$, $^{***}p< 0.001$, $^{****}p< 0.0001$, ns denotes not significant.

**d)** The normalized VEGFR1 expression levels at the plasma membrane are plotted against time. The mCherry intensity corresponding to VEGFR1 levels was normalized against the intensity of the membrane marker WGA-Alexa Fluor 633. The VEGFR1-mCherry constructs were transiently expressed in CHO cells and stimulated with 100ng/ml $VEGF_{165}$ (upper panel) or 100ng/ml $VEGF_{165}$ in the presence of 0.002 mM Razuprotafib (lower panel). Comparison between the groups were made using Student's t-test. Statistical significance was assessed using the following criteria: $^{*}p< 0.05$, $^{**}p< 0.01$, $^{***}p< 0.001$, $^{****}p< 0.0001$, ns denotes not significant

**e)** Bar graph representing relative mRNA levels of PTPRB determined from the quantitative PCR of the indicated cell lines.

**f)** Representative stepwise photobleaching traces of VEGFR1 clusters treated with 0.002mM Razuprotafib, plotted as mean intensity versus frame number. For mCherry bleaching, the laser power was set to 100%, and 400 frames were acquired every 4ms. 50 pre-bleach images and 5 post-bleach images were recorded. The arrow points to the distinct bleaching steps of the individual cluster.

**g)** Left is the representative whole-cell 2D STED images at the basal plane of HEK293T cells transiently expressing VEGFR1-mCherry (n=12-20 cells). VEGFR1 was stimulated with 0.002mM Razuprotafib. Right: a 4 µm x 4 µm representative region in the cell, showing a cluster considered for particle mobility analysis. The particle trajectory is shown in blue line, the start and end points marked in cyan and red, respectively.

**h)** Left is the same particle track, as in panel f, in a 1 µm x 1 µm region against a white background for clear visualization. The right panel shows the MSD of all individual particle tracks against time (from 0 to 0.06s). The individual tracks are shown in grey, and the mean is shown in red.

**b-f)** The plots are made using Graphpad prism 8.0.2. **g)** This plot is made using MATLAB R2026a.

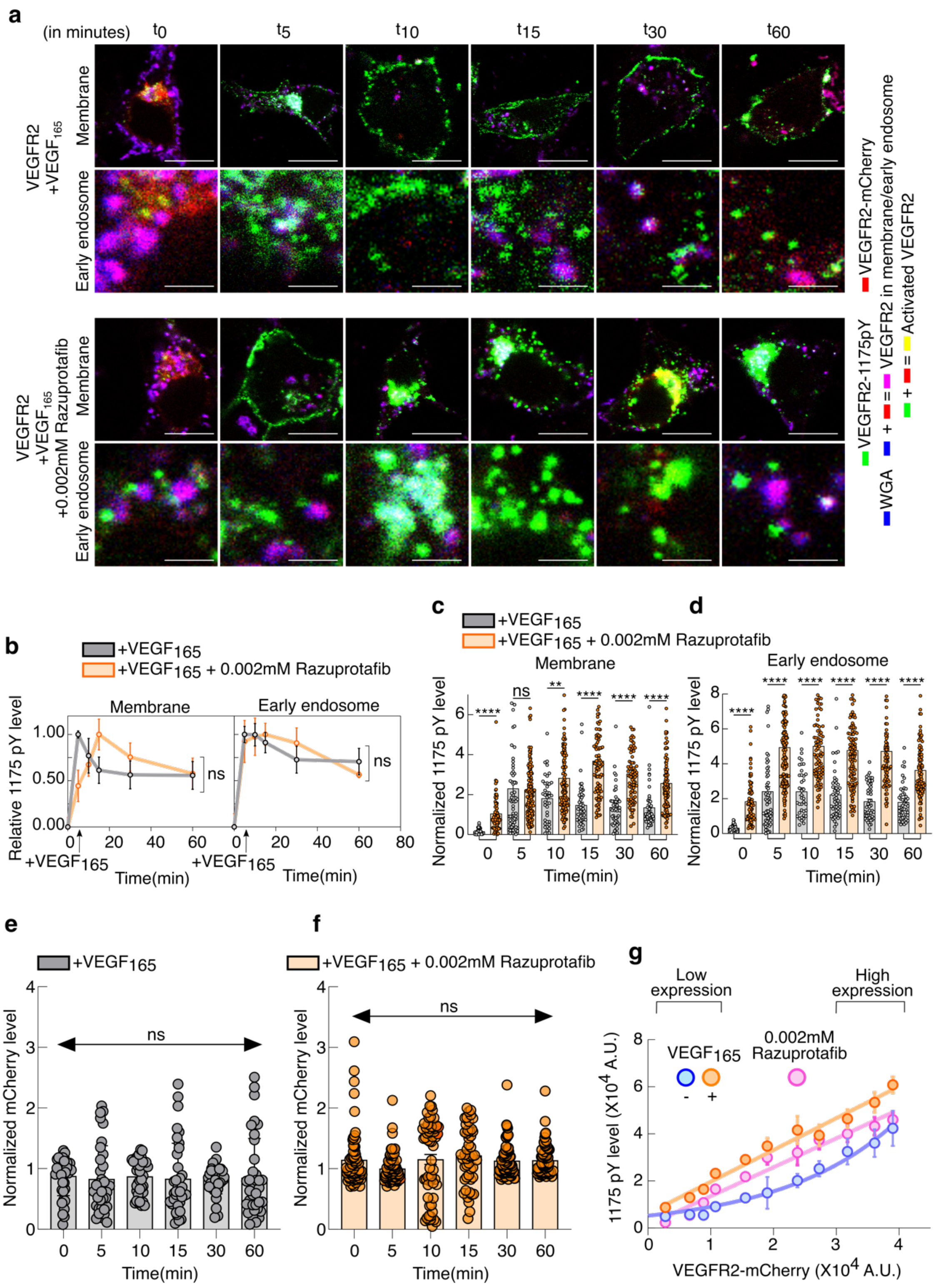


**Figure S7: Study of PTPRB on the rate of VEGFR2 autophosphorylation.**

**a)** Representative time-resolved confocal images of transiently VEGFR2 expressing HEK293T cells stimulation with 100ng/ml VEGF$_{165}$ (row 1 and 2) or 100ng/ml VEGF$_{165}$ in presence of 0.002mM Razuprotafib treatment (row 3 and 4). The images in rows 1 and 3 are of whole cells. Rows 2 and 4 are the images of the early endosome. The red channel ($\lambda_{ex}$ =552nm, $\lambda_{em}$ = 570-660nm), which denotes VEGFR2 expression. The green channel ($\lambda_{ex}$ = 488nm, $\lambda_{em}$ = 500-550nm) denotes Y1175 phosphorylation level (1175pY). The blue channel ($\lambda_{ex}$ = 638 nm and $\lambda_{em}$ = 650-725nm) denotes plasma membrane and early endosome labeled with WGA-tagged Alexa Fluor 633. The yellow and magenta represent the merged images. Cells expressing with mCherry intensity of 1-2 x $10^4$ AU at the plasma membrane were considered for this study. For rows 1 and 3, Scale bar=10 μm; For rows 2 and 4, Scale bar=1 μm.

**b)** The relative 1175pY level measured at the plasma membrane (left) or at the early-endosome is plotted against time. The HEK293T cell line transiently expressing VEGFR2 was activated with 100 ng/ml $VEGF_{165}$ (gray line) or 100 ng/ml $VEGF_{165}$ in the presence of 0.002mM Razuprotafib (orange line). Relative phosphorylation levels were determined by normalizing the background-corrected FITC: mCherry intensity ratio against the maximum. Each data point represents Mean ±SD from n=50-70 cells ($n_{total}$ = 450-495 cells). Comparison between the two groups at the 60-minute time point was made using Student's t-test. Statistical significance was assessed using the following criteria: $^{*}p< 0.05$, $^{**}p< 0.01$, $^{***}p< 0.001$, $^{****}p< 0.0001$, ns denotes not significant.

**c-d)** Bar plot of normalized 1175pY levels at the plasma membrane (panel c) or the early endosome (panel d) against time. The FITC intensity corresponding to 1175pY levels was normalized against theVEGFR2-mCherry intensity. Comparison between the two groups was made using Student's t-test. Statistical significance was assessed using the following criteria: $^{*}p< 0.05$, $^{**}p< 0.01$, $^{***}p< 0.001$, $^{****}p< 0.0001$, ns denotes not significant.

**e)** Bar plot of normalized mCherry intensity at the plasma membrane stimulated with 100 ng/ml $VEGF_{165}$ against time in HEK293T cells transiently expressing VEGFR2. The mCherry intensities were normalized against Alexa Fluor 633 intensity conjugated with membrane marker WGA. Comparison between the groups were made using Student's t-test. Statistical significance was assessed using the following criteria: $^{*}p< 0.05$, $^{**}p< 0.01$, $^{***}p< 0.001$, $^{****}p< 0.0001$, ns denotes not significant.

**f)** Bar plot of normalized mCherry intensity at the plasma membrane stimulated with 0.002mM Razuprotafib against time in HEK293T cells transiently expressing VEGFR2. The mCherry intensities were normalized against Alexa Fluor 633 intensity conjugated with the membrane marker WGA. Comparison between the groups was made using Student's t-test. Statistical significance was assessed using the following criteria: $^{*}p< 0.05$, $^{**}p< 0.01$, $^{***}p< 0.001$, $^{****}p< 0.0001$, ns denotes not significant.

**g)** Intensity of VEGFR2-mCherry at the plasma membrane of HEK293T is plotted against FITC intensity, denoting 1175pY level, n=150–200 cells. At each data point, cells were binned based on mCherry intensity. Each data point represents the mean ± SD from approximately 20–40 cells. The solid lines are guiding lines. VEGFR2 was activated either with 100 ng/ml $VEGF_{165}$ or with 0.002 mM Razuprotafib.

**b-g)** The plots are made using Graphpad prism 8.0.2.

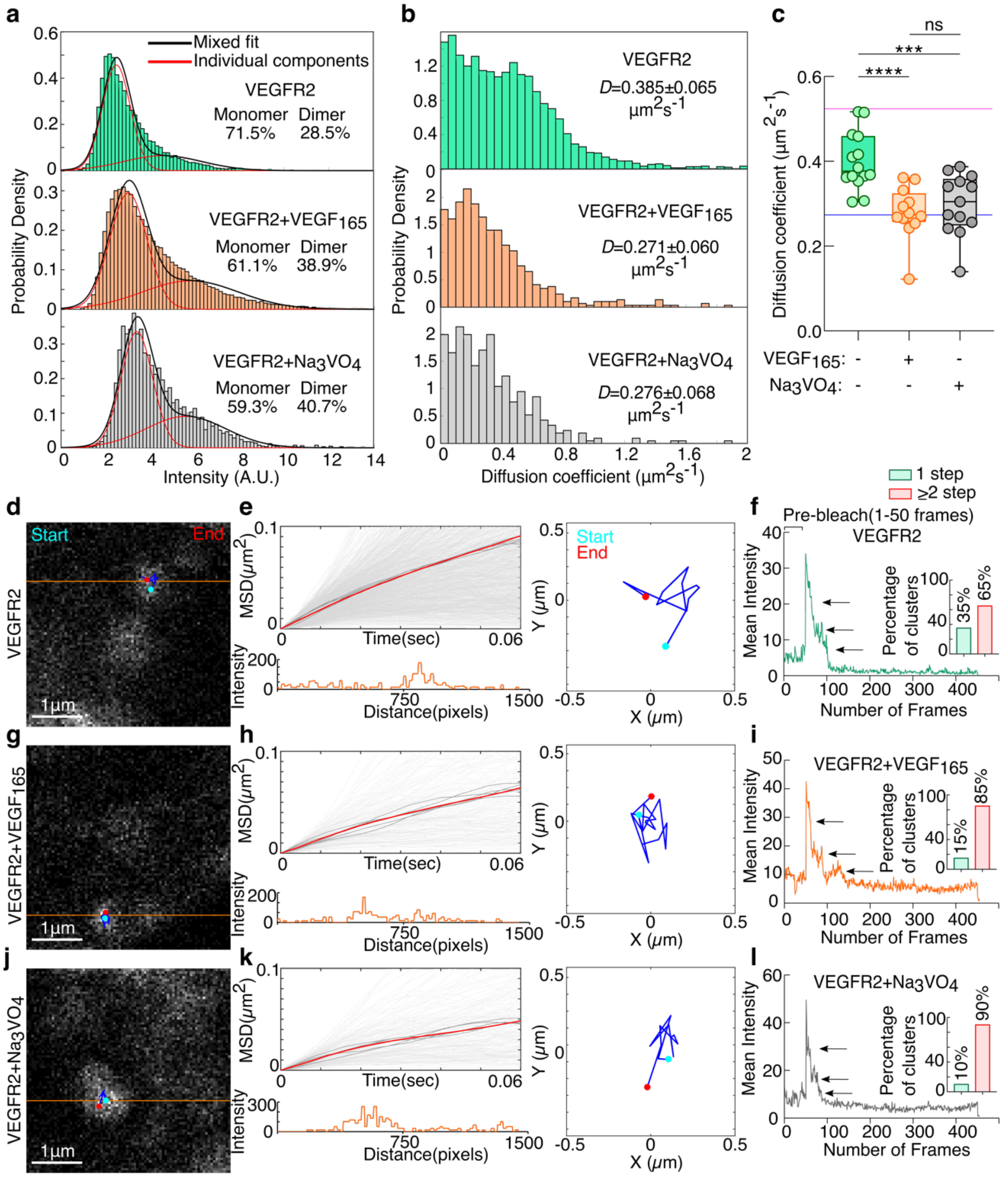


**Figure S8: Probing dimerization of VEGFR2 on the plasma membrane**

**a)** Histogram plot of intensity distributions of VEGFR2-mCherry against probability density. CHO cells transiently expressing VEGFR2 were activated with 100 ng/mL $VEGF_{165}$, or 0.1 mM $Na_3VO_4$. Each panel represents the intensity measured for all particles in n=10-20 cells. The red and black lines show single and mixed Gaussian fits, respectively. The percentages of monomer and dimer populations for each panel are reported.

**b)** Histogram distribution of diffusion coefficient for VEGFR2-mCherry either stimulated with 100ng/ml $VEGF_{165}$ or 0.1mM $Na_3VO_4$ is plotted against probability density. Each panel represents the diffusion coefficient measured for all valid tracks in n=10-20 cells. The diffusion coefficient represents the median ± SD.

**c)** The average diffusion coefficients of VEGFR2-mCherry measured in the absence of ligand (green) or after stimulating with 100 ng/ml $VEGF_{165}$ (orange) or in the presence of 0.1mM $Na_3VO_4$ (grey) from individual cells. Error bars represent mean ± SD from n = 10–20 cells. The solid pink line indicates the mean diffusion coefficient of VEGFR1-G83I (monomer control), and the blue solid line indicates the mean diffusion coefficient of VEGFR1-GPA (dimer control). Comparison between the two groups was made using Student's t-test. Statistical significance was assessed using the following criteria: $^{*}p< 0.05$, $^{**}p< 0.01$, $^{***}p< 0.001$, $^{****}p< 0.0001$, and ns denotes not significant.

**d-f); g-i); j-l)** are the representative images for particle mobility analysis of VEGFR2.

**d, g and j)** 4 µm x 4 µm representative region showing a cluster at the basal plane of CHO cells expressing VEGFR2-mCherry, considered for particle mobility analysis. The particle trajectory is shown in blue line, the start and end points marked in cyan and red, respectively.

**e, h and k)** Left is the mean Squared Displacement (MSD) of all individual particle tracks is plotted against time (from 0 to 0.06s). The individual tracks are shown in grey lines, and the mean is shown in a red line. The bottom panel shows the pixel intensity on the orange line drawn in the 4 µm x 4 µm representative 2D STED image. Right panel: same particle track in a 1 µm x 1 µm region, against a white background for clarity.

**f, i and l)** Representative stepwise photobleaching traces of VEGFR2 clusters, plotted as mean intensity versus frame number. For mCherry bleaching, the laser power was set to 100%, and 400 frames were acquired every 4ms. 50 pre-bleach images and 5 post-bleach images were recorded. The arrow points to the distinct bleaching steps of the individual cluster. In the inset, the bar plot shows the percentage of clusters exhibiting one-step (green bar) or two-step (red bar) bleaching (n=12-25 cells per condition).

**a, b, e, h, k)** Plots are made using MATLAB 2026a. **c, f, I, l)** plots are made using Graphpad Prism 8.0.2.

### References


1 Chakraborty, M. P. *et al.* Molecular basis of VEGFR1 autoinhibition at the plasma membrane. *Nat Commun* **15**, 1346, doi:10.1038/s41467-024-45499-2 (2024).

2 Schindelin, J. *et al.* Fiji: an open-source platform for biological-image analysis. *Nat Methods* **9**, 676-682, doi:10.1038/nmeth.2019 (2012).

3 Schmittgen, T. D. & Livak, K. J. Analyzing real-time PCR data by the comparative C(T) method. *Nat Protoc* **3**, 1101-1108, doi:10.1038/nprot.2008.73 (2008).

4 Arkhipov, A. *et al.* Architecture and membrane interactions of the EGF receptor. *Cell* **152**, 557-569, doi:10.1016/j.cell.2012.12.030 (2013).

5 Jaqaman, K. *et al.* Robust single-particle tracking in live-cell time-lapse sequences. *Nat Methods* **5**, 695-702, doi:10.1038/nmeth.1237 (2008).

6 Needham, S. R. *et al.* Measuring EGFR separations on cells with ~10 nm resolution via fluorophore localization imaging with photobleaching. *PLoS One* **8**, e62331, doi:10.1371/journal.pone.0062331 (2013).

7 Mudumbi, K. C. *et al.* Distinct interactions stabilize EGFR dimers and higher-order oligomers in cell membranes. *Cell Rep* **43**, 113603, doi:10.1016/j.celrep.2023.113603 (2024).

8 Calebiro, D. *et al.* Single-molecule analysis of fluorescently labeled G-protein-coupled receptors reveals complexes with distinct dynamics and organization. *Proc Natl Acad Sci U S A* **110**, 743-748, doi:10.1073/pnas.1205798110 (2013).

9 Huang, Y. *et al.* Molecular basis for multimerization in the activation of the epidermal growth factor receptor. *Elife* **5**, doi:10.7554/eLife.14107 (2016).

10 Cancer Genome Atlas Research, N. *et al.* The Cancer Genome Atlas Pan-Cancer analysis project. *Nat Genet* **45**, 1113-1120, doi:10.1038/ng.2764 (2013).

11 Goldman, M. J. *et al.* Visualizing and interpreting cancer genomics data via the Xena platform. *Nat Biotechnol* **38**, 675-678, doi:10.1038/s41587-020-0546-8 (2020).

12 Harris, C. R. *et al.* Array programming with NumPy. *Nature* **585**, 357-362, doi:10.1038/s41586-020-2649-2 (2020).

13 Edwards, N. J. *et al.* The CPTAC Data Portal: A Resource for Cancer Proteomics Research. *J Proteome Res* **14**, 2707-2713, doi:10.1021/pr501254j (2015).

14 Vasaikar, S. V., Straub, P., Wang, J. & Zhang, B. LinkedOmics: analyzing multi-omics data within and across 32 cancer types. *Nucleic Acids Res* **46**, D956-D963, doi:10.1093/nar/gkx1090 (2018).

15 Sondka, Z. *et al.* COSMIC: a curated database of somatic variants and clinical data for cancer. *Nucleic Acids Res* **52**, D1210-D1217, doi:10.1093/nar/gkad986 (2024).

16 Cerami, E. *et al.* The cBio cancer genomics portal: an open platform for exploring multidimensional cancer genomics data. *Cancer Discov* **2**, 401-404, doi:10.1158/2159-8290.CD-12-0095 (2012).

17 Huynh-Thu, V. A., Irrthum, A., Wehenkel, L. & Geurts, P. Inferring regulatory networks from expression data using tree-based methods. *PLoS One* **5**, doi:10.1371/journal.pone.0012776 (2010).
